# Evolution of a genome-architecture-encoded gene regulation system in trypanosomatids

**DOI:** 10.64898/2026.07.29.740908

**Authors:** Saurav Mallik, Meir Sylman, Moshe Kafri, Maayan Yoles, Bar Cohen, Dvir Dahary, Orna Dahan, Gerald F. Späth, Shulamit Michaeli, Yitzhak Pilpel

**Affiliations:** George S. Wise Faculty of Life Sciences, School of Neurobiology, Biochemistry, and Biophysics, Tel Aviv University, Tel Aviv 69978, Israel; Department of Molecular Genetics, Weizmann Institute of Science, Rehovot 7600001, Israel; The Mina and Everard Goodman Faculty of Life Sciences and Advanced and Nanotechnology Institute, Bar-Ilan University, Ramat-Gan 52900, Israel; Institut Pasteur, Université Paris Cité, INSERM U1347, Unité de Parasitologie moléculaire et Signalisation, Paris, France

**Author notes:** These authors contributed equally (Saurav Mallik, Meir Sylman). Corresponding authors: S.M., Y.P.

## Abstract

Transcriptional regulation of protein-coding genes is a hallmark of eukaryotic gene expression. Yet, a group of parasitic protists, trypanosomatids, appear to lack this capability. Here, we analyzed genomic, nascent transcriptomic, RNA polymerase occupancy and gene organization data to reconstruct the evolutionary origin and biological consequences of their unusual regulatory strategy. Across 59 Discoba protists, we show stepwise evolutionary erosion of conventional transcription regulation components in trypanosomatida lineage, including gene consolidation into polycistronic transcription units (PTUs), shortening of intra-PTU non-coding regions, and depletion of transcription factors and their enriched DNA-binding motifs. This transition was associated with near-constitutive expression of most genes, indicating broad loss of conditional gene expression. However, trypanosomatids retain some differential regulation at the PTU level, with >70% PTUs featuring significantly different nascent transcription than their neighbors or resident chromosomes. Moreover, gene expression is not uniform within PTUs: nascent transcription, translation efficiency, and protein abundance progressively decline with distance from the transcription start site. Consistent with this architecture-encoded regulatory logic, co-complex subunits and co-pathway enzymes preferentially occupy adjacent positions within PTUs despite each PTU’s overall functional heterogeneity. These findings reveal an evolutionary shift from gene-specific transcriptional regulation toward a regime where genome architecture becomes a regulator of gene expression.

## INTRODUCTION

All cellular functions, whether the assembly of protein complexes, the execution of metabolic pathways, responses to external perturbations, or cell division itself, rely on the precise spatiotemporal coordination of gene expression. Across the Tree of Life, this coordination is mediated by transcriptional and post-transcriptional regulation [1] – two shared processes driven by complex genetic and biomolecular interaction networks – that dictate when specific genes are activated, the temporal duration of their activity, stability of their mRNA transcripts, and their ultimate protein output.

Yet, a unique group of unicellular parasitic protists, namely trypanosomatids, are widely recognized to lack transcription regulation of protein-coding genes [2–8]. They feature densely packed, largely intronless genes, arranged in co-directional clusters known as polycistronic transcription units (PTUs), which resemble bacterial operons [9,10]. A single PTU can harbor anywhere from two to over a hundred genes that are often functionally unrelated [9,11,12] yet share a single transcription start and a stop site [2–7]. Gene order within these PTUs are remarkably conserved between distantly related trypanosomatids [13]. Each PTU is transcribed into a polycistronic pre-mRNA molecule, which is co-transcriptionally resolved into mature, monocistronic mRNAs via trans-splicing and polyadenylation [14]. Because this unique system fundamentally eliminates the independent transcriptional activation or differential expression of individual genes, gene expression in trypanosomatids is thought to rely almost exclusively on post-transcriptional mechanisms (modulating mRNA stability and translation) [2–7] or genomic copy number variations [15–17].

This radical departure from conventional transcriptional regulation raises fundamental evolutionary and regulatory questions. How did these polycistronic features originate? Through what sequence of events were individual promoters, transcription factors, and their binding motifs depleted in the trypanosomatid lineage? Is gene organization within these PTUs truly random, and is gene expression across them truly uniform? Finally, if these organisms cannot independently regulate individual gene expressions, how do they ensure the strict stoichiometric demands of multi-subunit complexes and metabolic pathways?

Here, we combined comparative genomics, data analysis of nascent transcriptomics and RNA polymerase occupancy, and functional gene organization to reconstruct how trypanosomatids transitioned from conventional gene-level transcriptional control to a genome-architecture-based regulatory regime. Across 59 Discoba species, we show that polycistronic gene arrangements arose through stepwise consolidation of genes into co-directional clusters, accompanied by shortening of intra-cluster non-coding regions and genome-wide depletion of transcription factors and their DNA-binding motifs. Although this transition was associated with a broad loss of conditional gene expression, we found that trypanosomatids retain some differential expression control at the PTU level. Moreover, PTUs are not uniform transcriptional units. Per-gene nascent transcription, translational efficiency, mRNA stability, and protein abundance progressively decline with its distance from the transcription start site. We further show that functionally coupled unrelated genes, including protein-complex subunits and metabolic-pathway enzymes, preferentially occupy adjacent positions within shared PTUs despite each PTU being overall functionally heterogeneous. Thus, gene-level differential regulation, to some extent, appears to be encoded in trypanosomatid’s genome architecture – order and adjacency of genes within PTUs. Altogether, our analysis places many previously documented findings about trypanosomatids into an evolutionary context and provides a systems-level understanding of their gene expression strategy.

## RESULTS

### Evolutionary diversification of Discoba protists and the origin of trypanosomatids

Several features of trypanosomatids’ unusual gene-expression strategy have been described previously [2–7], including polycistronic transcription, scarcity of conventional promoters and transcription factors, and extensive post-transcriptional regulation. However, these features have mostly been examined separately or within a limited set of model species. We sought to systematically examine the extent to which these features are conserved across trypanosomatids, how they differ from the corresponding features in their non-parasitic relatives, and when they emerged during Discoba evolution.

To this end, we assembled 59 high-quality protist proteomes (***Data S1***, ***Table S1***) and constructed a species tree. These proteomes spanned parasitic trypanosomatids [18], their closest free-living relatives predatory bodonoids [19,20], and representatives of diplonemid, euglenid, and other Discoba lineages. Following a systematic phylogenetic framework (***Methods***), we compared 973,660 proteins using all-versus-all BLAST [21] and clustered the resulting sequence-similarity graph into 48,018 distinct protein families [22] (***Data S2***). We then applied a maximum-likelihood approach [23] to a concatenated alignment of 154 highly conserved, nearly single-copy families to infer the species tree (**Figure 1A**, ***Data S3***). We rooted this tree using the aggregative multicellular amoeba *Acrasis kona*. This species is closely related to the free-living model protist *Naegleria* (phylum Heterolobosea) [24–26], which serves as an established outgroup to euglenids and diplonemids [27,28].

**Figure 1.**
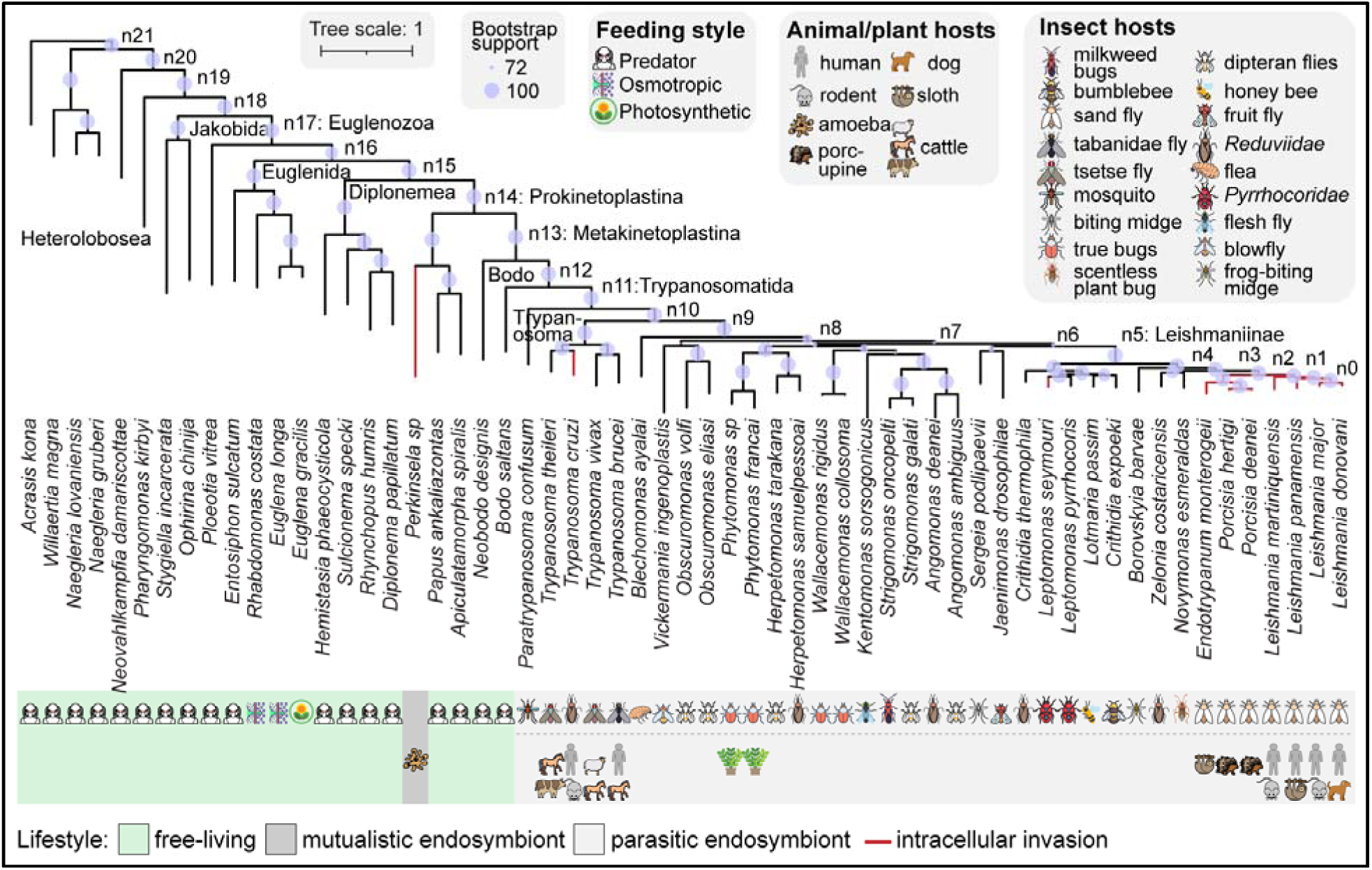
Evolutionary diversification of Euglenida, Diplonemida, and Kinetoplastida protists. A species tree representing the evolutionary divergence of 59 protist species with available ORFeome. The tree was inferred using maximum-likelihood methods from a concatenated alignment of 154 conserved, nearly single-copy protein families (***Methods***). Branch lengths represent the number of substitutions per 100 amino acids, and the size of the violet circles corresponds to bootstrap support values at internal nodes. Internal nodes relevant to downstream analyses are assigned arbitrar identifiers; clade names according to NCBI taxonomy [37] are provided when available. The species set includes free-living (green: predatory, photosynthetic, or osmotrophic), mutualistic endosymbionts (dar gray) and parasitic taxa (light gray). The lifestyle of each organism is indicated; for parasites, intracellular invasion capabilities are noted alongside their known insect vectors and vertebrate/plant hosts.

Overall, our species tree is consistent with recent reconstructions of protist phylogenies [18,29–31]. It supports multiple previously documented relationships, including (i) between monophyletic trypanosomatids and predatory bodonoids [19,20]; (ii) the sister relationships of *Wallacemonas* and *Strigomonas* [32]; (iii) the relatedness of *Vickermania* and *Jaenimonas* [18]; and (iv) the evolutionary splits between different *Trypanosoma* species [33]. Metakinetoplastina was confirmed to form two sister clades [27,29]: one containing trypanosomatida and eubodonida (*Bodo saltans*), and another containing neobodonida (*Neobodo designis*). Finally, Diplonemida, Euglenida, and Heterolobosea lineages emerged as monophyletic [27,29,34–36]. Collectively, these results confirm the reliability and robustness of our species tree and provide the necessary evolutionary scaffold for downstream comparative analyses.

### Stepwise erosion of gene-specific transcriptional regulation in the trypanosomatids

This species phylogeny allows us to systematically examine the evolution of trypanosomatid’s unique features. We started by examining polycistronic gene organizations [38,39] (**Figure 2A**). Transcription start and stop sites of PTUs are demarcated by chromatin marks [40] and the modified DNA base J [41], which remain unknown for most protists in our tree. Hence, we introduced a proxy: the ‘directon’, defined as a group of consecutive, co-linear genes on the same DNA strand (***Methods***, **Figure 2B**). In principle, a single directon can encompass one or more co-linear PTUs, and it enables a comparable genome-wide analysis across species even when chromatin- and base J-defined PTU annotations are unavailable. In reality, >80% PTUs in *L. major* and *T. brucei* genomes – the two organisms for which PTU annotations are available – harbor a single PTU (**Methods**, ***Figure S1***). This indicates that directons are reliable proxies for PTUs. By examining 11,477 genomes across the tree of life [42] (**Data S4**), we found that the average number of genes per directon (directon length) is ∼2 in plants, fungi, and animals (reflecting near-random strand assignment of genes) and 3-4 in prokaryotes (reflecting operon structures). Trypanosomatids stand out with average directon lengths of ∼30 (**Figure 2C**).

**Figure 2.**
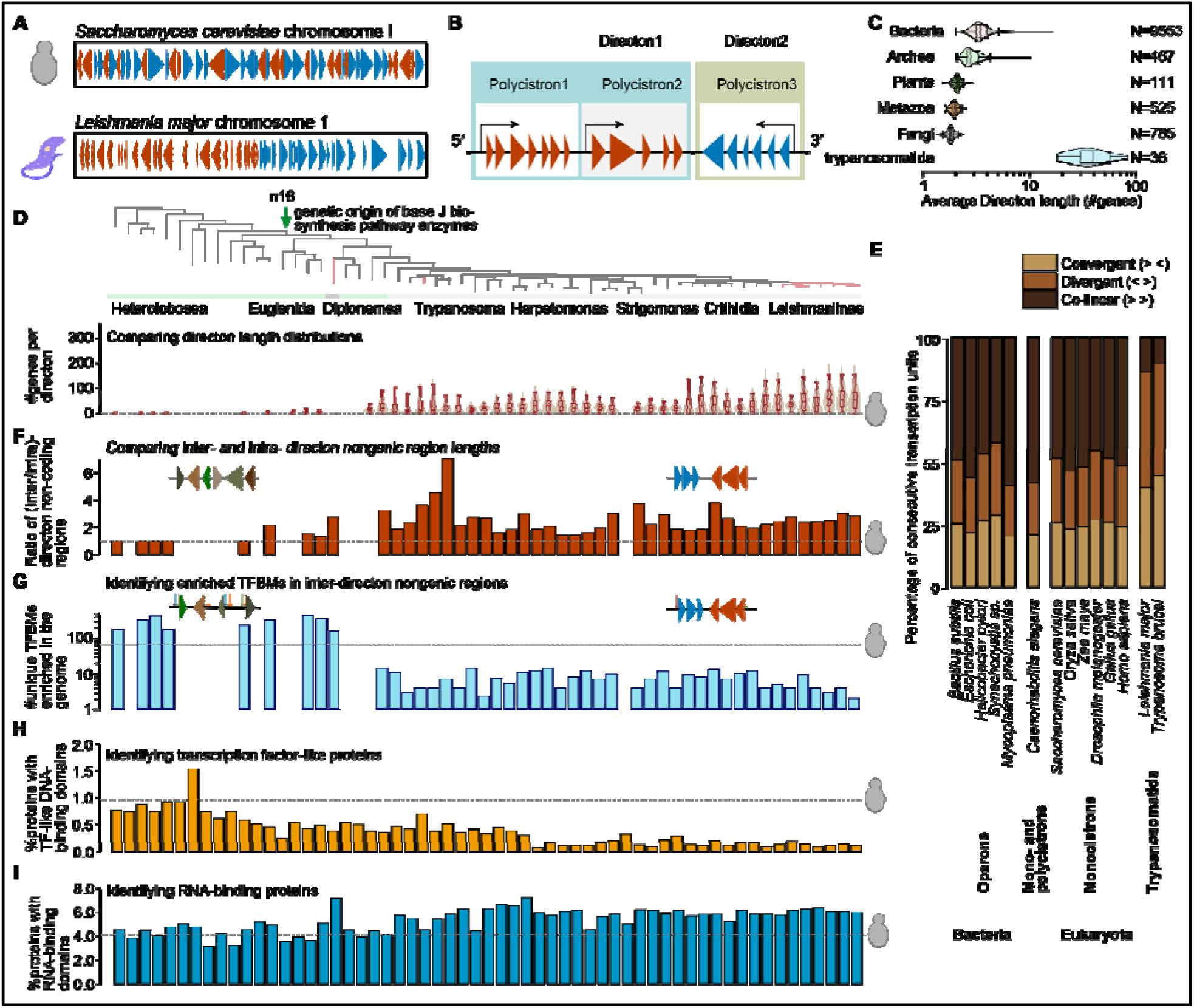
Systems-wide depletion of transcription regulation components in trypanosomatids. (**A**) Red and blue triangles illustrate the difference in gene orientations within a randomly selected 15-megabase region of budding yeast versus *L. major* chromosomes. (**B**) Schematic representation of a directon, which encompasses one or more co-linear PTUs. (**C**) Violin plots showing the distribution of the average directon lengths (#genes) per genome across the three domains of life. The number of species analyzed are mentioned. (**D-G**) Evolution of genomic features across the species tree. (**D**) The species tree, with major clades highlighted for reference, along with node n16 that marks the genetic origin of the enzymes in the base J biosynthesis pathway. Violin plots display the distribution of directon lengths across 47 protists with annotated genomes included in our species tree. Boxes and whiskers span the 25^th^-75^th^ and 5^th^-95^th^ percentiles, respectively. (**E**) Stacked bar plots show the percentage of consecutive transcription units per genome in convergent (yellow), divergent (light brown), and co-linear orientations (dark brown). The identity of a transcription unit varies between lineages; bacteria: operons, most eukaryotes: individual genes, *L. major* and *T. brucei*: PTUs, *C. elegans*, some transcriptional units are PTUs, others are individual genes. (**F**) Same as **D**; bar plots represent the ratio of total inter-directon versus intra-directon non-coding region lengths in the genome. (**G**) Same as **D**; bar plots represent the count of unique transcription factor binding motifs (TFBMs) enriched in inter-directon non-coding regions. (**H-I**) Evolution of proteome features across the species tree. (**H**) Same as **D**; bar plots represent the percentage of proteins in the proteome harboring DNA-binding domains of known transcription factor activity. (**I**) Same as **D**; bar plots represent the percentage of genes in the genome harboring RNA-binding domains involved in mRNA stability, localization, and translational control activity. Panels **F-I** are aligned to the species tree shown in panel **D**; the horizontal dotted lines reflect the corresponding value for the budding yeast genome. Genomic features include data for taxa with high-quality genomes; proteome features include data for all species.

We reasoned that if this contrast is captured within our species tree, tracing these gene arrangements across genomes would allow us to delineate the stepwise evolution of the polycistronic transcription system. Results show that early-diverging Heterolobosea and Jakobida protists exhibit ∼2 genes/directon (***Data S5***). A departure from this trend occurs at node n16 in our species tree: its two descendant lineages, the euglenids and diplonemids, exhibit average directon lengths of ∼4 and ∼6, respectively (**Figure 2D**, ***Figure S2***). Indeed, extant members of both clades exhibit polycistronic transcription [43–45]. Strikingly, this departure coincides with the evolutionary emergence of the base J biosynthesis pathway. In present-day *L. major* cells, base J, an epigenetic modification of thymine, is synthesized on the DNA in two enzymatic steps [46]. First, thymidine hydroxylases oxidize methyl groups on specific thymines, producing HOMedU. Next, a β-glucosyltransferase converts HOMedU into base J by adding a glucose molecule. Our phylogenetic analysis traced the origin of both enzymes to node n16. This coincidence suggests that the evolutionary innovation of base J may have enabled the expansion of polycistronic architectures, allowing their regulated transcriptional termination. Notably, base J usage is not uniform across Euglenozoa. In the Euglena *gracilis* genome, base J is more uniformly distributed and likely plays a minor role in transcription termination [43,47]; in trypanosomatids base J is clustered in telomeric regions and defines PTU boundaries.

The co-linear gene clusters have increased dramatically in the Discoba lineage leading to *Leishmania* (**Figure 2D**). The directon length distribution of all Euglenozoa species exhibit pronounced heavy-tails, *i.e.*, unusually long directons in their genomes. This heavy-tailedness is absent in Heterolobosea and Jakobida protists (***Figure S2***). The longest directon in the genome increases from 58 in *Euglena gracilis* to 99 in *Diplonema papillatum*, and 325 in *L. major*, indicating increased consolidation of genes into polycistronic arrangements. This consolidation is captured in ***Figure S3***, which plots the genomic fraction of PTUs containing ≥X and ≤Y genes. By shifting these parameters (X from 10 to 80, and Y from 2 to 10), we map how, along the Discoba lineage toward *Leishmania*, longer directons progressively take over the genomes.

Surprisingly, PTUs appear to be non-randomly organized within trypanosomatid chromosomes. In most eukaryotes, transcriptional units are individual genes, and these genes are randomly oriented within chromosomes. As a result, ∼25%, ∼25%, and ∼50% of their consecutive gene pairs are found in divergent, convergent, and co-linear orientations, respectively (**Figure 2E**). Similar trends exist in bacteria at operon level, and also in roundworm *Caenorhabditis elegans*, where some transcriptional units are polycistronic and others are monocistronic [48,49]. By contrast, trypanosomatid chromosomes are enriched for divergent and convergent PTUs (∼45% and ∼40%) and depleted for co-linear ones (∼15%). These biases likely minimize transcriptional readthrough, *i.e.* RNA polymerase II leakage, from one PTU into the next. Such leakage is known to occur in trypanosomatids, both naturally [50], and upon base J disruption [50–52].

We reasoned that incorporation of formerly monocistronic genes into PTUs should have reduced the length of intergenic regions separating co-linear genes, because these regions no longer harbor individual promoters. We show this erosion in **Figure 2F**, with the caveat that our intergenic region annotations also include 5′ and 3′ UTRs (***Methods***). In budding yeast, inter- and intra-directon intergenic regions are roughly equal in length (ratio ≈1). Early-diverging Heterolobosea and Jakobida maintain this trend. However, in the lineage toward *Leishmania*, this ratio steadily climbs, eventually hovering around ∼2 in the trypanosomatida genomes. Importantly, since these intergenic regions also include untranslated regions, the true reality of promoter loss could be more drastic than depicted in **Figure 2F**.

It is generally accepted in trypanosomatid biology that their genomes largely lack transcription factors [8,53–56]. Because polycistronic architectures arose gradually along the lineage leading to *Leishmania*, we asked whether this transition was mirrored by a corresponding evolutionary depletion of transcription factors (TFs) and their binding motifs (TFBMs) from the genome.

To address this question, we retrieved all known eukaryotic TFBMs from the JASPAR database [57,58] and used the MEME suite [59,60] to scan inter-directon non-coding regions. Since TFBMs are short (6-15 base pairs) and can easily occur by chance in megabase-scale genomes, we filtered strictly for those significantly overrepresented against a background model of randomly shuffled sequences (***Methods***). This conservative approach identified 65 unique enriched motifs in the monocistronic budding yeast genome. The early-diverging Heterolobosea and Jakobida genomes, being 3-4 times larger than yeast, harbor hundreds of unique enriched TFBMs. Strikingly though, this repertoire virtually collapses in the trypanosomatids, dropping to <10 TFBMs per genome in *Leishmania* (**Figure 2G**, ***Data S6***). Furthermore, TFBM loss exceeds what would be expected due to the reduction of transcription units per genome (***Figure S3***). We note that this analysis is limited by the current knowledge of known TFs in model eukaryotes, and is expected to lack lineage-specific TFs that may exist in these genomes.

To trace evolutionary dynamics of genes encoding for transcription factors themselves, we compiled 65 Pfam-annotated [61] DNA-binding domains associated with TF activity in eukaryotes [62–65] (***Table S2***) and quantified their representation across the species tree (***Methods***). Whereas the budding yeast genome encodes 58 proteins (∼1% of all protein coding genes in the genome) harboring at least one such domain, early-diverging Heterolobosea and Jakobida, with genomes 3-4 times larger than yeast, typically encode ∼100 (∼0.7% of all protein coding genes). This TF repertoire sharply declines along the trypanosomatids lineage, reaching ∼8 in *Leishmania* (∼0.1% of all protein coding genes; **Figure 2H**).

Finally, since differential gene regulation in trypanosomatids is thought to rely almost exclusively on post-transcriptional mechanisms [2–7], we compiled 190 Pfam-annotated [61] RNA-binding domains associated with eukaryotic proteins known to modulate mRNA stability, localization, and translational control [66–68] (***Table S3***, ***Methods***). Across our species tree, early-diverging Heterolobosea and Jakobida display ∼800 RNA-binding proteins (RBPs) per proteome (∼4% of protein coding genes); this matches the proportion found in budding yeast (**Figure 2I**). The absolute number of RBPs decreases to ∼430 in the *Leishmania* proteomes due to overall genome streamlining, yet their proportion in the proteome increases to ∼6%. This increased RBP proportion is consistent with their prominent role in trypanosomatid gene regulation [69,70].

Taken together, using directons as proxies for PTUs, our findings highlight the gradual rise of the unique, largely unregulated polycistronic transcription system in trypanosomatids. The evolutionary onset of these directional gene clusters coincides with the emergence of base J biosynthetic enzymes. As genes were consolidated into PTUs, the shortening of intergenic regions indicates a systematic erosion of individual promoters. The depletion of transcription factors and their binding motifs, combined with the rise of RNA-binding proteins reflect a global transition towards largely unregulated transcription and increased dependency on post-transcriptional regulation. Finally, a genome-wide bias against co-linear PTUs indicates an evolutionary selection against RNA Pol-II leakage from one PTU to the next.

### Differential expression of PTUs is retained in trypanosomatids amidst the loss of conditional gene expression regulation

We asked how the evolutionary transition described above contributed to a shift away from conventional transcription regulation in trypanosomatids. We reasoned that the latter acts on a transcriptional unit through two complementary mechanisms: a binary decision of whether it is active or inactive (an on/off *switch*) and the ability to independently fine-tune its transcriptional output (a *knob*, **Figure 3A**). The identity of the transcription unit in this analysis varies across lineages: individual genes in most eukaryotes, operons in prokaryotes, and PTUs in trypanosomatids.

**Figure 3.**
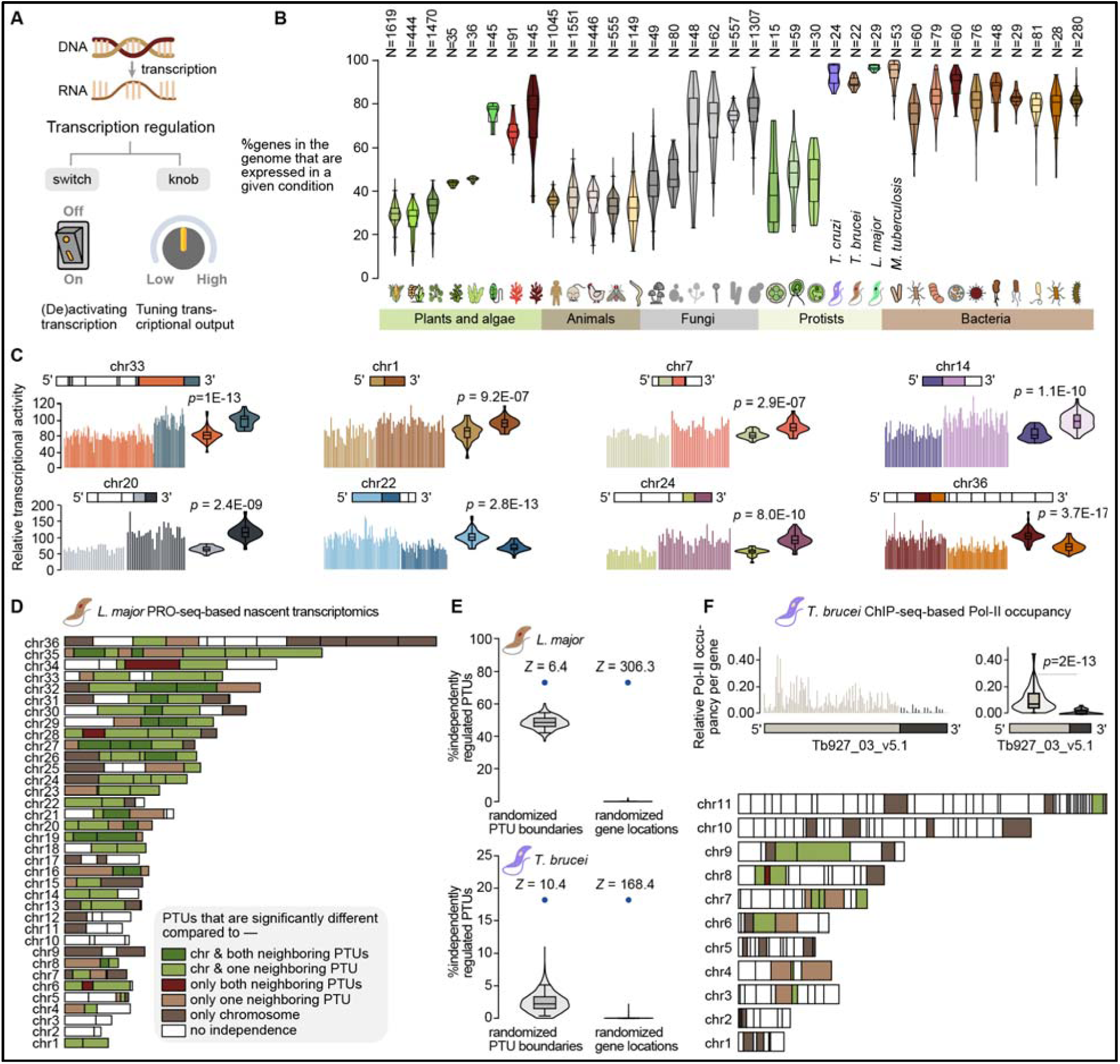
Trypanosomatids appear to differentially express most of their PTUs. (**A**) A schematic representation of the *switch* and the *knob* components of conventional transcription regulation. (**B**) The presence of the *switch* component across the Tree of Life. For each organism, the violin plots represent the percentage of genes in the genome that are transcriptionally active (transcript per million or *TPM* >5) across multiple RNA-seq datasets. The number of datasets for each taxa are mentioned. (**C**) Bar plots comparing the per-gene nascent transcription rates for eight consecutive PTU pairs (highlighted in different colors) from various chromosomes of *L. major* genome. The same data i also shown as violin plots, where each violin corresponds to a PTU. The *p*-value was derived from the Mann-Whitney U test. (**D**) A map of the *L. major* genome: PTUs harboring ≥3 genes are represented as rectangles. The colors represent whether a PTU is transcribed at a significantly different rate from its neighbor(s) and/or its corresponding chromosome. The transcription rates were obtained from a PRO-seq-based nascent transcriptomics study [73]. (**E**) We compared the frequency of PTUs displaying a *knob* to a null model. The latter was derived iteratively (N = 1000) by shuffling PTU boundaries while keeping gene order fixed (left violin) and by shuffling gene order while keeping PTU boundaries fixed (right violin). The Z-scores represent how many standard deviations the original data (blue circle) deviates from the mean of this null model. (**F**) Same as **D**, for *T. brucei*, using relative per-gene Pol-II occupancy derived from ChIP-seq-based footprints [77] (see ***Figure S5***).

To determine whether trypanosomatids have lost these regulatory capabilities, we assessed the retention of the *switch* component across the tree of life. We analyzed 10,607 RNA-seq datasets spanning 38 model organisms, focusing on healthy wild-type samples and excluding gene knockouts, mutations, disease states, and stress treatments where possible (***Table S4***). For each organism and dataset, we calculated the fraction of annotated genes that were detectably expressed, using a threshold of >5 transcripts per million (TPM).

Our results indicate a broad range across species, featuring species in which only about a third of all genes are active in any given condition, to species in which most genes are active in each condition. For example, close to the high end of the scale, 75-80% of bacterial genes are expressed in each condition, yet only 20-40% of plant and animal genes are active at any given condition (**Figure 3B**). This contrast is consistent with bacteria’s operon-based genome architecture, where activation of a single operon turns on multiple genes simultaneously. Across eukaryotes, increased organismal complexity typically corresponds with more conditionally activated genes (more *switches*). For example, as plant complexity increases from unicellular algae to non-vascular moss to vascular crops like rice and corn, the percentage of active genes per condition steadily drops from ∼70% to ∼45% to ∼30%. Trypanosomatids stand out as outliers: 93-95% genes are constitutively activated, suggesting that they have largely lost conditional gene expression through the *switch* mechanism.

While the vast majority of trypanosomatid genes are constitutively activated, their exact expression level may still be differentially regulated at the PTU-level, in a *knob* like fashion. We reasoned that if a PTU’s transcriptional output differs significantly from that of its genomic neighbors or its resident chromosome, this would provide evidence for a regulatory *knob*. To systematically identify those PTUs, we split our directons into PTUs using histone acetylation marks [71] and then analyzed a PRO-seq-based [72] nascent transcriptomics dataset [73] for *L. major* and compared per-gene transcription rates (**Data S7**, ***Methods***). Because PRO-seq captures actively engaged RNA polymerases (Pol-II) incorporating biotinylated nucleotides during a nuclear run-on assay, it directly measures nascent transcriptional activity [73]. In this study, during logarithmic growth of *L. major* cells at 25°C in M199+ medium, nascent transcription levels were found to vary across PTU boundaries of the same chromosome [73]. This striking pattern is depicted in **Figure 3C**. Adjacent PTUs on various *L. major* chromosomes exhibit marked differences in per-gene nascent transcription output, amidst relative constancy within their respective boundaries. This trend is reminiscent of differential regulation of adjacent bacterial operons [74–76]. However, these trends could also originate from subchromosomal amplifications, yet only in the unlikely event in which they precisely overlap PTU boundaries. While such amplification events were not explicitly accounted for, they are primarily triggered by environmental stressors, which doesn’t apply to these experiments.

We scaled this analysis to systematically compare every PTU against its neighbors and resident chromosomes. We found that PTU-level transcriptional *knobs* are prevalent in the *L. major* genome (**Figure 3D**). Specifically, 125 PTUs (73%, out of 171 containing ≥3 genes with rate data) exhibit differential regulation signatures: they showed significantly different per-gene nascent transcription outputs relative to at least one neighbor or their resident chromosome. The dynamic range of this differential regulation – the maximum observed nascent transcription output difference between PTUs in the same chromosome – appears to be around 3.0-fold. We confirmed the statistical significance of our result against two null models: one by randomly shuffling genes within chromosomes while keeping PTU boundaries fixed, and another by shuffling PTU boundaries while keeping gene order fixed (***Methods***). Neither of these negative controls reproduced the frequency of differentially regulated PTUs observed in the actual genome (one-sample right-tailed Z-test; Z = 6.4 and 306.3, respectively, **Figure 3E**).

We asked whether a fundamentally different experimental approach, namely Pol-II footprints obtained from ChIP-seq experiments [77], would reproduce similar trends for *T. brucei*. While PRO-seq specifically captures transcriptionally active Pol-IIs at nucleotide resolution [73], ChIP-seq-based footprints capture a snapshot of Pol-II engagements, also at nucleotide resolution, but cannot determine if they are actively elongating or stalled. Using these footprint data for *T. brucei* cells cultured in HMI-11 medium at 37°C, we calculated relative Pol-II occupancy per gene (**Data S8**, **Methods**). Here too though, these occupancies differed significantly across PTU boundaries, exemplified between two adjacent PTUs on chromosome 3 (Mann-Whitney U-test *p* = 2E−13, **Data S9**, **Figure 3F**, see ***Figure S4*** for additional examples). Again, the frequency of PTUs with significantly different occupancies (relative to ≥1 neighbor or chromosome) observed in the actual genome was not reproduced in our two null models (Z = 10.4 and 168.4, **Figure 3E**).

Together, these analyses separate two fundamental aspects of transcription regulation that are often conflated in the literature. Those include the capability for (i) conditional activation of a gene and (ii) differential regulation of its expression level. The former is nearly eliminated in trypanosomatids; though the latter appears to have been largely retained. In the absence of abundant conventional transcription factors, PTU-level differential regulation is likely mediated through chromatin remodeling [78–80]. Our results are consistent with a recent nascent transcriptomics study in *T. cruzi*, which revealed a group of ‘core’ PTUs that are associated with open chromatin marks. These PTUs contain housekeeping genes, and are transcribed at a significantly higher rate than a group of ‘disruptive’ PTUs containing mostly virulence genes [81].

### Positional expression gradients are embedded within polycistronic architectures

Different cellular functions require distinct levels of gene expression [82,83] and protein abundances [84]. However, a typical trypanosomatid PTU harbors tens to hundreds of largely functionally unrelated genes [9,11,12] that are transcribed at identical levels. While this discrepancy is widely thought to be reconciled by post-transcriptional regulation, we asked whether a gene’s precise sequential order in the PTU carries specific regulatory significance. Such a trend is not without precedent. Previous studies have shown that during heat shock, a gradient of gene expression fold change (between two conditions) appears along the trypanosomatid PTUs: genes close to TSS are downregulated, while those far away are upregulated [85]. This happens because heat shock likely halts transcription initiation, while elongation continues; due to the finite speed of transcription, genes far away from the TSS are transcribed later [85].

However, this transient, stress-induced phenomenon does not address whether a gene’s physical position influences its steady-state mRNA level in a single condition. Strikingly, we observe that these levels (obtained by RNA-seq [86]) progressively decrease as a function of the gene index, *i.e.* its position along the PTU. Hence, genes closer to the TSS are expressed on average two-fold more than the distal ones. This decay is illustrated in **Figure 4A** for representative PTUs in *L. major* and *T. brucei* genomes. To examine this effect on a genome-wide scale, we constructed a meta-polycistron profile. For *L. major* cells grown at 26°C, we normalized the steady-state mRNA level of each gene [86] (TPM values) to the mean expression of its resident PTU, and aligned all PTUs containing ≥10 genes relative to their first gene (***Methods***). This analysis revealed a continuous decrease in mean normalized steady-state mRNA levels as the gene index increases (*R^2^* = 0.66, *p* = 1.8E−24, **Figure 4B**). Similar results were derived for *T. brucei* cells in culture [87] (*R^2^* = 0.63, *p* = 2.3E−21, **Figure 4C**; see ***Figure S5*** for additional conditions). Accordingly, we also found that both in *L. major* and *T. brucei*, highly and lowly expressed genes in a given condition tend to locate near and far from the TSS, respectively (***Figure S6***).

**Figure 4.**
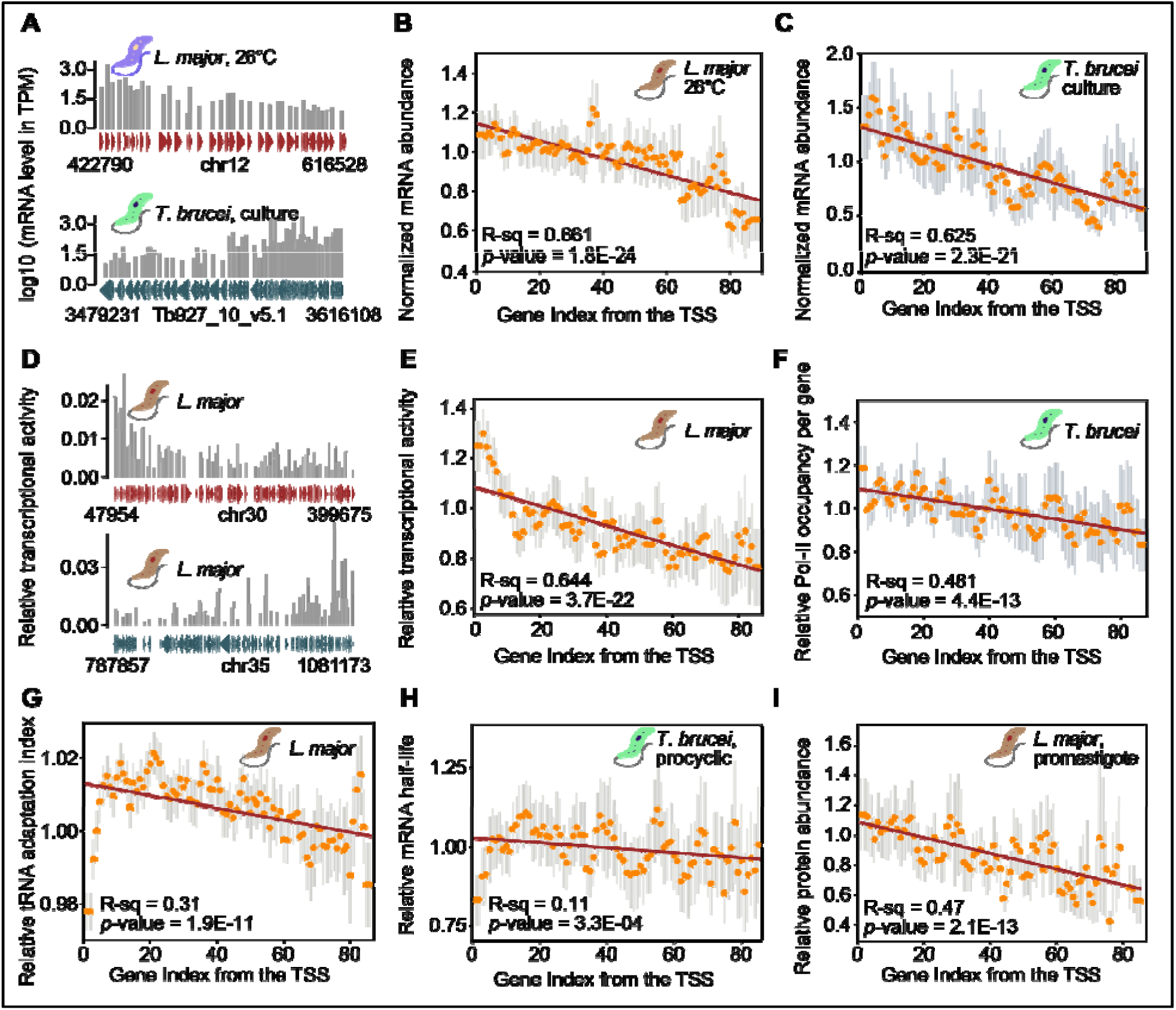
A regulation gradient is embedded in the polycistronic structures. (**A**) For individual PTUs in *L. major* (26°C, red) and *T. brucei* (culture growth, blue), gray vertical bars represent log10-transformed steady-state mRNA levels (in transcripts per million, or *TPM*) for individual genes. Genes are represented as triangles pointing in the direction of transcription; chromosome identifiers and genomic coordinates are mentioned. (**B**) Average mRNA levels of *L. major* cells grown at 26°C [86] are aligned with the index of their respective first genes. The orange points and gray vertical lines represent the per-gene average mRNA level, normalized to the average mRNA level of its PTU, and the respective standard deviations. The red line represents the linear regression; the corresponding *R^2^* and *p*-value are mentioned. (**C**) Same as B, for *T. brucei* cells in culture [87]. (**D**) Same as A, for per-gene relative transcriptional activity (relative Pol-II occupancy per gene per kilobase, derived from PRO-seq-based nascent transcriptomics data [73]), for two example *L. major* PTUs. (**E**) Same as B, for per-gene nascent transcription output in *L. major* cells. (**F**) Same as B, for relative Pol-II occupancy per gene, derived from ChIP-seq experiments in *T. brucei* cells [88]. (**G**) Same as B, for tRNA adaptation index per gene, for the *L. major* genome. (**H**) Same as B, for mRNA half-life per gene, for *T. brucei* cells in the procyclic stage [87]. (**I**) Same as **B**, for protein abundance level in the *L. major* cells in the promastigote stage [89].

We reasoned that this gradient may originate from premature transcription elongation termination [90], which is common in bacteria [91,92], eukaryotes [93–95] and also in trypanosomatids [96–99]. Specifically, localized enrichment of the modified DNA base J, the histone variant H3.V, or the protein phosphatase PP1 are known to actively promote Pol-II termination before the end of a PTU [96–99]. However, whether these events are pervasive enough to generate a continuous, genome-wide negative gradient of nascent transcription rates remains unknown. To this end, we analyzed a PRO-seq-based nascent transcription dataset [73]. Specifically, across two representative PTUs on *L. major* chromosomes 30 and 35, genes positioned closer to the TSS are transcribed more than the distal ones (**Figure 4D**). A meta-polycistron profile establishes this principle on a genome-scale: the mean normalized nascent transcription rate per gene drops steadily as the gene index increases (*R^2^* = 0.64, *p* = 3.7E−22, **Figure 4E**). The regression curve suggests an approximate ∼0.21% loss of actively elongating Pol-IIs per kilobase in *L. major*, which is comparable to ∼0.35% measured in aging human liver [95]. Similar results were obtained for *T. brucei* using per-gene relative Pol-II occupancy data derived from ChIP-seq experiments [88] (*R^2^* = 0.48, *p* = 4.4E−13, **Figure 4F**).

Having established expression level gradients along the trypanosomatid PTUs, we turned to post-transcriptional regulation. This decline in mRNA expression with distance from the TSS can be physiological, originating from Pol-II terminations of along PTUs. Yet it could also have an evolutionary origin, whereby genes with longer mRNA half-life are preferentially positioned closer to PTU boundaries. We thus examined gene properties that are less likely to be affected by the act of transcription itself. In particular we examined the tRNA adaptation index (tAI) of genes along PTUs, which is a measure of their codon adaptation to the tRNA pool of the species, and mRNA half-life [100] (***Methods***, ***Data S10***). Mirroring the transcriptional data, tAI steadily decreases with increasing gene index (*R^2^* = 0.31, *p* = 1.9E−11, **Figure 4G**, also see ***Figure S7***). Notably, the first 3-4 genes in this meta-polycistron profile exhibit a significantly lower baseline tAI before the values peak and begin their descent. This is reminiscent of the established translational ramp at the mRNA level, wherein the initial 30-50 codons are translated with low efficiency [101,102].

We observed a corresponding, though milder, positional decay in mRNA stability for *T. brucei* [87]: mRNA half-life decreases with gene index following a similar initial ‘ramp’ consisting of the first 4-5 genes being less stable (*R^2^* = 0.11, *p* = 2.3E−04, **Figure 4H**). This correspondence between low codon to tRNA adaptation and mRNA half-life is consistent with recent literature that ascribes an mRNA de-stabilizing effect to lack of codon optimality [103]. Finally, we asked whether these transcriptional and post-transcriptional gradients manifest at the protein level too. By evaluating three independent *L. major* proteomics datasets [89,104,105], we confirmed that mean normalized per-gene protein abundance also follows this negative positional gradient (*R^2^* = 0.47, *p* = 2.1E−13, **Figure 4I**, also see ***Figure S8***).

Together, these analyses reveal that trypanosomatid PTUs are not uniform transcriptional units. Instead, they contain multiple positional gradients: genes closer to the TSS tend to show higher nascent transcription, better codon-tRNA adaptation, slightly greater mRNA stability, and higher protein abundance than distal genes. These gradients likely provide a vital gene-level differential regulation in these protists. It further suggests that evolution may have exploited the order of genes within PTUs as a regulatory parameter, whereby genome architecture itself induces differential gene regulation.

### Functionally coupled genes are non-randomly arranged within trypanosomatid PTUs

Having established that trypanosomatid PTUs differ in transcriptional output and contain positional gradients in gene expression, we next asked whether these regulatory properties are reflected in present-day gene organization along PTUs. Since physical proximity within a PTU should induce similar gene expression, genes requiring tight stoichiometric synchrony should be biased toward physical proximity within the same PTU. Trypanosomatid PTUs are generally thought to contain largely functionally unrelated genes [9,11,12]. This notion, however, does not exclude the possibility that small subsets of genes requiring tight synchrony are non-randomly positioned within PTUs.

We focused on two classes of biochemical systems expected to impose relatively strong constraints on component abundance: multisubunit protein complexes [106–108] and multi-enzyme metabolic pathways [109–111]. We compiled enzyme/subunit compositions for annotated eukaryotic complexes (obtained from Complex Portal [112]) and metabolic pathways (obtained from MetaCyc [113]) and mapped their orthologs in the *L. major* genome (***Methods***, ***Data S11***), identifying 1,609 unique subunit genes and 994 unique enzyme genes. For each subunit/enzyme, we considered only one *L. major* ortholog with the highest sequence similarity, to avoid a false positive adjacency signal originating from gene arrays – groups of near-identical protein-coding genes organized in tandem repeats. We found that only a small fraction of unrelated gene pairs, 3.9% (750 pairs) for complex subunits and 4.1% (370 pairs) for pathway enzymes, reside within the same PTU. This confirms that PTUs are not generally organized as functionally coherent operons [9,11,12]. Yet, these co-occurrence rates were far higher than expected from random gene pairs (one-sample right-tailed Z-test against a null distribution of 10,000 random pairs; Z = 45.0 and 37.6; **Figure 5A**), indicating that a small group of genes belonging to the same complex/pathway are selectively enriched within shared PTUs. Indeed, paralogous genes are pervasive in protein complexes [114] and metabolic pathways [115] and since they were excluded from our analysis, we expect the true enrichment of functionally related genes in the *L. major* genome to be significantly higher. Such an analysis, however, awaits the availability of high-quality genome annotations.

**Figure 5.**
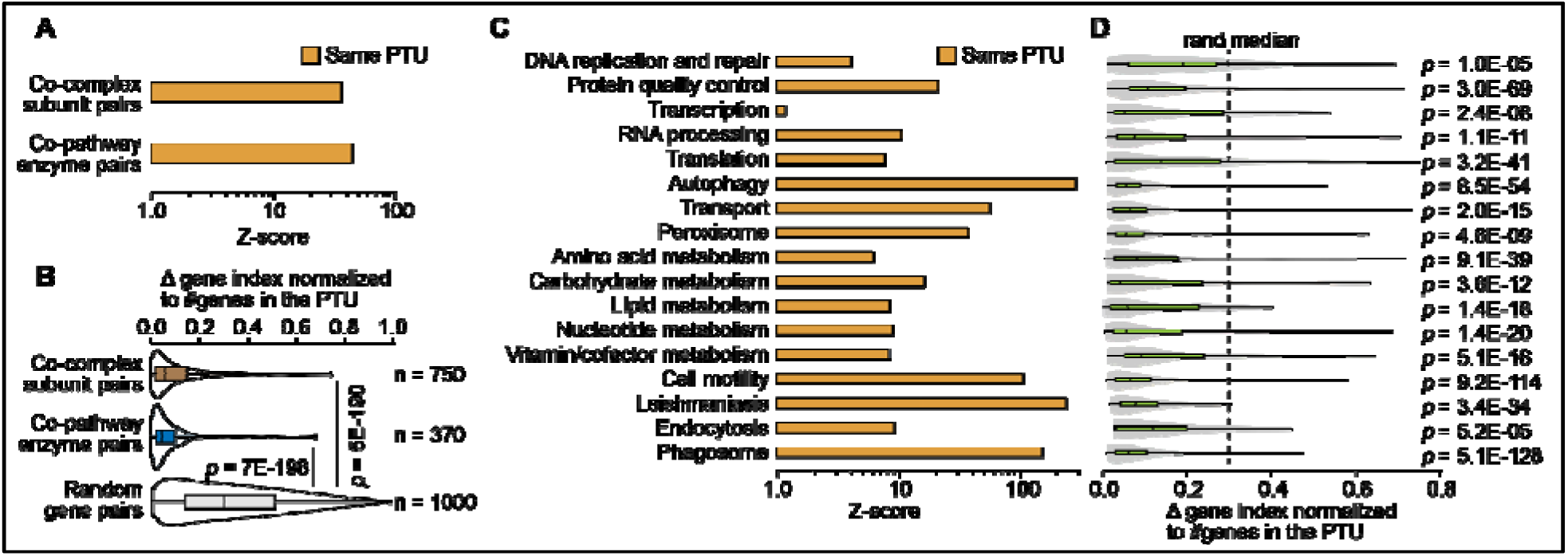
Gene organization in trypanosomatid PTUs. (**A**) Bar plots represent enrichment of gene pairs encoding subunits of the same complex or enzymes of the same metabolic pathway within *L. major* PTUs. The enrichment is quantified in terms of a Z-score against a background of 10,000 random gene pairs. (**B**) Violin plots comparing the gene index deviations (normalized by the total number of genes in the PTU) for gene pairs encoding subunits of the same complex (yellow) or enzymes of the same metabolic pathway (blue) within the same PTU. The gray violin represents the same for 1000 random gene pairs within the same PTUs. (**C**) Same as **A**, for KEGG-annotated broader gene functional classes. (**D**) Same as **B**, for the same broader gene functional classes. The vertical dashed line represents the median for random pairs. The *p*-values were derived from pairwise Mann-Whitney U-tests against this random distribution.

We then asked whether functionally coupled gene pairs that share a PTU are randomly distributed within it or preferentially positioned near one another. We investigated the spatial arrangement of the 750 co-complex and 370 co-pathway gene pairs that indeed reside on the same PTU. We found that their gene indices were significantly more adjacent than expected by random chance (two-sample Mann Whitney U test, *p* = 5E−190 and *p* = 7E−198, **Figure 5B**), indicating a strong tendency to be adjacent within the PTU.

Finally, we asked whether these biases are unique to biochemical systems requiring strict stoichiometric synchrony (complexes and pathways), or whether they also govern broader functional classes like DNA replication, autophagy, and RNA processing. While these broader classes require distinct gene expression and protein abundance levels [82,84], they are not expected to require strict stoichiometric synchrony among their components. To test this, we classified the *L. major* proteome using KEGG [116] into several major functional classes (**Figure 5C-D**, ***Data S12***). Each functional class encompasses hundreds of proteins. We found that gene pairs from the same broad functional category tend to co-occur within the same PTU more frequently than expected by chance (**Figure 5C**). Moreover, when these pairs do localize within the same PTU, they maintain a significant bias toward being adjacent (**Figure 5D**). Notably, strongest enrichments, both for co-occurrence within the same PTU and gene adjacency within PTUs, are observed for genes related to autophagy, leishmaniasis, cell motility, and phagosome that are central to the parasites’ survival within macrophages. Of note, a weaker, albeit significant co-localization tendency also exists at the chromosome level (***Figure S9***).

Taken together, these findings suggest present-day trypanosomatid gene organization is globally heterogeneous but locally structured: functional relationships are not sufficient to define entire PTUs, yet PTU membership and within-PTU gene orders are locally biased to favor small groups of functionally coupled genes. This suggests that a unique genome organization has evolved in trypanosomatids’ exploiting their unique gene expression strategy. Specifically, functionally coupled genes are biased to co-occur within physical proximity within the same PTU, presumably to ensure their similar transcriptional outputs and protein abundances.

## DISCUSSION

Our results support a model in which trypanosomatids underwent an evolutionary transition from conventional gene-specific transcriptional regulation toward a regulatory regime encoded partly in genome architecture. Comparative analyses across Discoba protists indicate that this transition involved the progressive expansion of directional gene clusters, shortening of intra-cluster intergenic regions, and depletion of transcription factors and enriched DNA-binding motifs. These changes are associated with a broad reduction in gene-level conditional switching, resulting in near-constitutive expression of most genes. However, these protists did not eliminate differential regulation altogether, which is retained at the level of PTUs. Additionally, individual gene regulation is encoded in the gene order within PTUs: nascent transcription, tRNA adaptation, and protein abundance decline with distance from the transcription start site. Thus, the apparent loss of conventional promoter-based regulation of individual genes is replaced by an alternative regulatory regime where a gene’s PTU-affiliation and within-PTU positioning influences its expression level.

Our results point towards a remarkable phase of genome architectural evolution in early Euglenozoans when individual genes started being consolidated into polycistronic clusters. The evolutionary onset of this process appears to coincide with the emergence of base J biosynthesis pathway. Base J a modified thymine that epigenetically marks polycistron boundaries [41]. Which genetic mechanism could have enabled this consolidation? One plausible route is retrotransposon- or integrase-mediated insertion [117,118]. Most of these insertions lack proper promoters and cannot be expressed unless they land within an existing PTU. Retrotransposons and repetitive elements are indeed pervasive in present-day *Diplonema* and *Euglena* genomes [28,119–121]. Notably, these PTUs are arranged within trypanosomatid chromosomes avoiding co-linear orientations. This may help minimize transcriptional readthrough when RNA polymerase II leaks from one PTU into the next.

Our findings regarding differential regulation of PTUs are consistent with recent GRO-seq-based nascent transcriptomic analyses in *T. cruzi* [81]. This study established a fundamental connection between a PTU’s gene content and its transcriptional output, identifying highly expressed ‘core’ PTUs containing housekeeping genes alongside lowly expressed ‘disruptive’ PTUs containing mostly virulence genes. Crucially, elevated transcription in ‘core’ PTUs correlated with open chromatin marks [81], suggesting that in the absence of conventional transcription factor activity, differential regulation of PTUs is likely mediated by chromatin remodeling [78–80].

Trypanosomatids appear to differentially regulate the transcriptional output of individual PTUs while simultaneously subjecting them to an intrinsic gradient of gene expression. We found a nascent transcription rate gradient within PTUs; this gradient likely emerges from premature elongation termination of RNA Pol-II molecules. Similar expression gradients have been discovered for polycistronic transcription of nematode [122] and mammalian [123] mitochondrial genomes, as well as for mobile integron systems in bacteria [124,125]. It appears that in trypanosomatids, evolution has extended this nascent transcription gradient to the protein level by also arranging genes of better tRNA adaptation closer to the TSS. The modest decline in mRNA half-life appears to mirror the trend obtained for tAI, and reflects the fact that codon optimality is a determinant of mRNA stability [103]. However, there is another possibility that during transcription, some information is embedded within the mRNA that determines its stability [126,127]. According to such a model, the Pol-II profile along PTU may have an effect on the decline in stability of the transcript. These positional gradients should not be interpreted as biologically independent effects: steady-state mRNA abundance integrates nascent transcription and mRNA decay [127], while tRNA adaptation of codon usage can influence both mRNA stability and translational efficiency [103]. Consequently, the protein-abundance gradient likely arises as a downstream consequence of the other gradients.

Furthermore, trypanosomatids frequently use tandem repeats, extrachromosomal amplicons, and chromosomal aneuploidy to boost gene expression, particularly under stress conditions [15–17]. Under both transcriptional and gene-dosage regimes, physical proximity warrants similar expression levels. Indeed, gene pairs required to maintain strict mRNA-level stoichiometric synchrony are biased to co-localize within the same PTUs and maintain adjacent positioning. Though, globally, functional couplings between genes are not sufficient to define entire PTUs.

Notably, only a small fraction (∼4%) of all co-complex subunit pairs or co-pathway enzyme pairs reside within the same PTU. This modest fraction was obtained by disregarding tandem gene duplicates and they support the conclusion that trypanosomatid PTUs are not equivalent to bacterial operons, nor are they generally organized as coherent functional modules. Nevertheless, PTU membership and within-PTU gene order are locally biased for small sets of functionally coupled genes. The biological relevance of this modest degree of colocalization is supported by our previous work on protein complexes, which showed that strict 1:1 stoichiometry is required for only ∼5% of human co-complex subunit pairs, particularly those that are mutually stabilized in the oligomeric state and tend to misfold in isolation [107,128]. Having most subunits of a complex requiring strict stoichiometric synchrony or else misfolding is likely a huge burden on the protein quality control machinery [128,129] and such trends are typically avoided in evolution [130]. Thus, the local colocalization observed here likely reflects a biologically meaningful constraint acting on a limited subset of functionally coupled genes.

In summary, our analysis systematically characterizes trypanosomatids’ gene regulation regime, while placing many previously documented findings systematically into a unified evolutionary and regulatory framework. The emergence of long PTUs was accompanied by erosion of conventional promoter- and transcription factor-based individual gene control, but not by a complete loss of transcriptional regulation. Instead, trypanosomatids appear to have shifted toward a regime in which differential regulation has been rescaled at the PTU-level, and also partially encoded in the genome architecture. These regulatory features, together with gene copy-number variations and post-transcriptional regulation, shape gene expression and protein abundance. More broadly, this work illustrates how the evolutionary loss of one regulatory regime can be accompanied by the emergence of another, in which genome architecture itself becomes a new regulatory parameter.

## Acknowledgments

We thank all members of the Michaeli, Späth, Pilpel, and Mallik lab for helpful discussions. This work was supported by the ERC 2022-SYG DECOLeishRN grant to S.M. (Bar-Ilan), G.F.S., and Y.P., and by the Minerva Center for Live Emulation of Evolution in the lab to Y.P.

## Author contributions

All authors participated in conceiving, conceptualizing the project and analyzing the data, S.M., M.S., M.K., B.C., and M.Y. performed the analyses, S.M. supervised the project, prepared the figures, and wrote the first draft. Y.P., acquired funding and supervised the study. All authors discussed and commented on the results and edited versions of the paper.

## Data and Materials availability

All data used or generated in this study are provided as Supplementary Materials.

## Materials and Methods

### 1. Taxon sampling, genome/proteome sources, and quality assessment

For this study, we curated a taxon set of publicly available protist proteomes based on three primary criteria. First, to trace the evolutionary origins of polycistrons, the dataset needed to encompass organisms both with and without directional gene clusters. The former includes all Euglenozoa, while the latter includes other members of the Discoba supergroup. We restricted our outgroup species to free-living Heterolobosea and Jakobida protists. Second, to avoid taxonomic bias, we aimed to sample uniformly across this evolutionary range by excluding highly redundant closely related taxa. For example, although high-quality annotated genomes are available for dozens of *Leishmania* and *Trypanosoma* species, we selected only four representatives for each. Third, when multiple closely related genomes were available, we prioritized those with the highest quality in terms of both contiguous genome assembly and gene annotation.

Applying these criteria, we sampled 59 protist species with publicly available predicted proteomes (**Table S1**, **Data S1**). This included 37 trypanosomatids and the predatory Bodo saltans, for which fully sequenced and annotated genomes were available from previously described resources [18], as well as 21 outgroup species representing additional Discoba lineages [24,27,29,131–133]. For 47 species, genome assemblies and gene annotations were available and were used for genome-architecture analyses. For the remaining 12 species, genome assemblies were unavailable; these taxa were therefore used exclusively for proteome-based analyses (e.g., transcription factor proteins per proteome), and not for genome-based analyses (e.g., enriched transcription factor binding sites).

Genome assembly completeness was taken into account by prioritizing genomes with chromosome-level or supercontig-level assembly, when possible. Proteome completeness was assessed using BUSCO v6.1.0 [134] against either the Eukaryota dataset, comprising 255 conserved single-copy orthologs, or the Euglenozoa dataset, comprising 130 conserved single-copy orthologs, depending on the taxonomic classification of each species. The average BUSCO completeness score was 93.4% for trypanosomatids and 72.0% for outgroup species. Lower BUSCO scores in several outgroups may reflect both technical and biological factors, including incomplete transcriptome-derived proteomes, limited lineage-specific representation in available BUSCO datasets, gene loss in highly adapted or parasitic protists, and difficulties in orthology assignment among rapidly evolving protist genomes [135–137]. Species sources, data types, and BUSCO scores are provided in **Table S1**.

### 2. Protein family inference and species-tree reconstruction

Protein families were inferred from the 59 predicted proteomes using OrthoFinder v2.5.5 [22] with multiple-sequence-alignment-based tree inference enabled (-M msa -S mmseqs -A muscle -I 1.5). In total, 973,660 annotated protein sequences were compared by all-versus-all BLASTp-based [21] sequence-similarity search. To reduce spurious similarity links caused by short or partial alignments, query-subject pairs were excluded when the aligned region covered <30% of either the query or subject protein sequence. The filtered similarity graph was then clustered using an inflation parameter of 1.5, yielding 48,018 protein families (**Data S2**).

For species-tree reconstruction, we selected 154 conserved, nearly single-copy protein families from the OrthoFinder output (**Data S3**). Protein sequences from each family were aligned using T-Coffee in M-Coffee mode [138], which integrates alignments generated by PCMA [139], MAFFT [140], ClustalW [141], POA [142], Muscle [143], and tcoffee [138] (-mode mcoffee - gapopen 20 -gapext 5). Individual family alignments were trimmed with trimAL [144] to remove poorly aligned and gap-rich columns (-gt 0.6 -cons 50). The resulting curated alignments were concatenated into a supermatrix.

To reduce potential phylogenetic artifacts caused by compositional heterogeneity, the 20% most compositionally heterogeneous sites were removed from the concatenated alignment [145,146]. The best-fitting amino-acid substitution model was selected using ModelFinder according to the Bayesian Information Criterion. The selected model was LG+F+I+G4 (-LnL = 2461441.24, BIC = 4924164.64). Maximum-likelihood phylogenetic inference was then performed using IQ-TREE v1.6.12 [23] under the LG+F+I+G4 model, with 1,000 ultrafast bootstrap replicates (-m LG+F+I+G4 -bb 1000). The resulting species tree was rooted using the heterolobosean amoeba Acrasis kona as the outgroup. Protein family data, along with the list of 154 nearly-single-copy families, are provided in **Data S1**.

### 3. Genome-architecture annotation: directons and PTUs

Genome-architecture analyses were performed for the 47 species in our dataset for which both genome assemblies and gene annotations were available (**Table S1**). We defined a directon as a maximal set of consecutive genes encoded on the same DNA strand. Directons were annotated using in-house Python scripts that scanned each genome annotation, ordered genes by genomic coordinate within each chromosome, scaffold, or contig, and assigned consecutive co-oriented genes to the same directon. Both protein-coding genes and annotated non-coding RNA genes were included in directon annotation.

Genome assembly contiguity varied across species (**Data S5**). Chromosome-scale assemblies were available for several model organisms, including *Leishmania major*, *Trypanosoma brucei*, and *Euglena gracilis*, whereas many other genomes were assembled at contig or supercontig scale. For contig- or supercontig-level assemblies, directons located at the 5′ or 3′ end of each contig were excluded from directon-length analyses, because such terminal directons may represent truncated fragments of longer directional gene clusters. Internal directons were retained. Only one trypanosomatid, *Angomonas deanei*, was excluded from genomic analysis as the quality of its contig-level genome assembly was too fragmented.

For each genome, we calculated directon length as the number of genes contained within each directon. To compare directon organization across major domains of life, we additionally analyzed 11,383 chromosome-scale genome assemblies from Ensembl [42], comprising 9,496 bacterial, 467 archaeal, 111 plant, 525 metazoan, and 784 fungal genomes (**Data S4**). Directons were annotated in each genome as described above, and the mean number of genes per directon was calculated for each species.

To examine changes in the shape of directon-length distributions along the lineage toward trypanosomatids, we used in-house Python scripts to fit directon-length distributions from selected representative genomes to geometric, power-law, and lognormal models. Model fits were compared using the Bayesian Information Criterion (BIC), and the model with the lowest BIC was considered the best-fitting distribution.

For analyses requiring experimentally defined PTU boundaries, directons were further partitioned into PTUs in *L. major* and *T. brucei*. In *L. major*, PTU boundary definitions were obtained from a previous study [71] that characterized histone acetylation marks that demarcate RNA polymerase II transcription start regions. In *T. brucei*, PTU boundary definitions were obtained from a previous study [88] that characterized RNA polymerase II transcription start regions by tracing its footprint density peaks on the genome. These species-specific PTU annotations are provided in **Data S9**. Strikingly, we found that the majority of directons in *L. major* and *T. brucei* genomes harbor only one PTU (***Figure S1***).

To compare non-coding sequence organization across genomes, we classified intergenic intervals as either intra-directon or inter-directon. Intra-directon intervals were defined as non-coding regions between consecutive genes within the same directon. Inter-directon intervals were defined as non-coding regions separating adjacent directons. For each genome, we summed the total length of all intra-directon and inter-directon intergenic intervals and calculated the ratio of inter-directon to intra-directon non-coding sequence. Because these annotation-derived intervals include untranslated regions as well as potential promoter-associated sequences, this analysis was used as an indirect measure of changes in non-coding architecture rather than as a direct measurement of promoter loss.

Additionally, operon annotations for Bacillus subtilis, Escherichia coli, Helicobacter pylori, Synechocystis sp., Mycoplasma pneumoniae, and Caenorhabditis elegans were obtained from the OperonDB [147] database.

### 4. Annotation of regulatory protein repertoires

We annotated two classes of regulatory proteins across the 59 protist proteomes: (i) proteins harboring DNA-binding domains that typically associate with transcription factor activity and (ii) proteins harboring RNA-binding domains that typically associate with post-transcriptional regulation. These annotations were based on conserved Pfam domains [61] and were therefore used to quantify domain repertoires across species, rather than to assign experimentally validated regulatory functions to individual proteins.

To define transcription-factor-associated DNA-binding domains, we first compiled experimentally characterized eukaryotic transcription factors from AnimalTFDB [62], PlantTFDB [65], P2TF [63], and TFClass [64]. For each transcription factor, we extracted its Pfam-annotated domain architecture [61] and retained domains corresponding to DNA-binding activity. Basal transcription factors were excluded from this analysis. This procedure yielded 65 unique Pfam DNA-binding domains associated with eukaryotic transcription factors (**Table S2**).

To define RNA-binding domains associated with post-transcriptional regulation, we used experimentally characterized RNA-binding proteins previously implicated in mRNA stability, localization, or translational control [66–68]. Pfam-annotated domains were extracted from these proteins as described above, yielding 190 unique RNA-binding domains (**Table S3**).

The 65 transcription-factor-associated DNA-binding domains and 190 RNA-binding domains were then searched against all 59 predicted proteomes using HMMER [148] and the corresponding Pfam hidden Markov model profiles. Hits were retained if they passed an e-value cutoff of <10^−5^. For each proteome, we counted the number of proteins containing at least one domain from each domain set and calculated the corresponding fraction of the annotated proteome. Proteins containing multiple domains from the same set were counted once for that regulatory class.

### 5. Transcription-factor binding motif enrichment analysis

To assess whether the depletion of transcription-factor-associated DNA-binding domains was mirrored by depletion of their enriched DNA-binding sequence motifs, we analyzed inter-directon non-coding regions from the 47 species with available genome assemblies and annotations. Inter-directon regions were defined as non-coding intervals separating adjacent directons, and were used as the sequence space most likely to contain promoter-associated regulatory elements. For each genome, the corresponding inter-directon sequences were extracted and used as the foreground sequence set.

As a background control, each foreground sequence set was randomly shuffled to preserve the overall nucleotide composition while disrupting motif organization. We retrieved 4,572 position frequency matrices representing known eukaryotic transcription-factor binding motifs from the JASPAR database [57,58]. Motif enrichment in each genome was tested using the SEA package from the MEME suite [59,60], comparing motif occurrences in the original inter-directon sequences against the shuffled background sequences. Only those motifs with an e-value < 0.01 were considered enriched. For each genome, we counted the number of unique enriched transcription-factor binding motifs (**Data S6**).

Because the number of directons varies substantially across species, motif counts were also normalized by the number of directons per genome where indicated. This analysis was used to quantify the repertoire of enriched known eukaryotic transcription-factor binding motifs in promoter-proximal inter-directon regions. It does not exclude the existence of lineage-specific motifs or regulatory elements absent from current motif databases.

### 6. Cross-kingdom RNA-seq analysis of gene activity

To compare the extent of gene-level transcriptional activity across diverse organisms, we analyzed 10,607 RNA-seq-derived gene expression datasets spanning 38 model organisms across bacteria, archaea, fungi, plants, animals, and protists. Processed expression matrices were obtained from comparative transcriptomics resources, including Bgee [83], Expression Atlas [149], Fungi.guru [150], Bacteria.guru [151], Protist.guru [152], and others [153–159]. Only datasets reporting gene-level expression values as transcripts per million (TPM) were included (**Table S3**).

Where possible, we focused on healthy wild-type samples and excluded datasets corresponding to genetic perturbations, mutant strains, disease states, and overt stress treatments. For unicellular organisms, individual datasets typically represented distinct growth conditions or life-cycle stages. For multicellular organisms, datasets typically represent organs, tissues, sub-tissues, cell types, or developmental contexts.

For each dataset, genes with expression >5 TPM were classified as transcriptionally active. We then calculated the fraction of annotated genes that were active in each dataset. This metric was used as a standardized proxy for the fraction of the genome transcriptionally active in a given biological condition. Although this approach does not capture all forms of condition-specific regulation, it enables a uniform comparison of gene activity across organisms and transcriptomic resources.

### 7. PTU-level nascent transcription and RNA polymerase occupancy analyses

To quantify PTU-level transcriptional output in *Leishmania major*, we analyzed a previously published PRO-seq-based nascent transcriptomic dataset [73]. Processed read counts mapped to each annotated gene, including coding and untranslated regions, were length-normalized and scaled across the transcriptome to obtain per-gene TPM values. These values were then assigned to the corresponding *L. major* PTUs.

For each PTU, we compared the distribution of per-gene nascent transcription values with three local genomic contexts: the upstream neighboring PTU, the downstream neighboring PTU, and genes on the rest of the chromosome. Comparisons were performed using two-sided Mann-Whitney U tests followed by false-discovery-rate correction. A PTU was classified as having differential transcriptional regulation if its per-gene transcription distribution differed significantly from at least one neighboring PTU or from its resident chromosome.

To determine whether the observed frequency of PTUs with differential transcriptional output could arise from genome organization alone, we generated two sets of randomized genomes using in-house Python scripts. In the first null model, gene order was shuffled within each chromosome while PTU boundaries were kept fixed. In the second null model, PTU boundaries were shuffled within each chromosome while preserving gene order and the number of PTUs per chromosome. For each null model, 1,000 randomized genomes were generated and analyzed using the same PTU-level comparison pipeline as the original genome.

We performed an analogous analysis for *Trypanosoma brucei* using previously published ChIP-seq-based RNA polymerase II footprint data mapped at nucleotide resolution [88]. Footprint density within each annotated gene, including coding and untranslated regions, was length-normalized and scaled across the genome to obtain relative Pol-II occupancy per gene. Per-gene occupancy values were assigned to *T. brucei* PTUs and analyzed using the same PTU-level comparison and randomization framework described above. Because Pol-II ChIP-seq measures polymerase occupancy rather than active nucleotide incorporation, this analysis was interpreted as an independent proxy for PTU-level transcriptional engagement rather than as a direct measure of nascent transcription rate.

### 8. Positional gradients along PTUs

To test whether gene position within PTUs is associated with gene-expression output, we analyzed multiple gene-level measurements of gene/mRNA/protein properties and projected them onto experimentally annotated *L. major* and *T. brucei* PTUs. These measurements included steady-state mRNA abundance, nascent transcription, RNA polymerase II occupancy, tRNA adaptation index, mRNA half-life, and protein abundance.

Steady-state mRNA levels for *L. major* promastigotes grown at 26°C and 35°C were obtained from RNA-seq-derived TPM values [86]. Steady-state mRNA levels for *T. brucei* cells in culture, procyclic, and bloodstream stages were obtained from published RNA-seq datasets [87]. Gene-level nascent transcription values from PRO-seq [73] in *L. major* and relative Pol-II occupancy values from ChIP-seq [88] in *T. brucei* were processed as described above and analyzed using the same positional framework.

For each dataset, gene-level values were assigned to their corresponding PTUs. To compare genes across PTUs with different overall expression levels, each gene’s value was normalized by the mean value of all genes in its resident PTU. We refer to this PTU-normalized value as the normalized gene-level output. PTUs containing fewer than 10 annotated genes were excluded from positional-gradient analyses. The remaining PTUs were aligned by gene index relative to the first gene downstream of the experimentally annotated transcription start site. For each gene-index position, we calculated the mean and standard deviation of the normalized values across all aligned PTUs containing a measured value at that position, generating a meta-polycistron profile for each molecular measurement.

To estimate predicted translational efficiency, we calculated the tRNA adaptation index (tAI) for annotated protein-coding genes in *L. major* and *T. brucei*. tRNA genes were identified using tRNAscan-SE [160]. The tAI values were calculated using an in-house MATLAB implementation based on fixed wobble-penalty weights derived from yeast (**Data S10**).

For mRNA stability, we analyzed a genome-scale dataset of mRNA half-lives measured in the *T. brucei* procyclic stage [87]. These half-lives were originally inferred from transcription inhibition followed by RNA-seq across multiple time points. For protein abundance, we analyzed three independent *L. major* proteomics datasets [89,104,105]. The mRNA half-life and protein-abundance values were assigned to PTUs, normalized within PTUs, and incorporated into meta-polycistron profiles as described above.

For each meta-polycistron profile, we evaluated the relationship between normalized gene-level output and gene index using linear regression across all gene-index positions.

### 9. Functional gene-organization analyses

To test whether functionally coupled genes are non-randomly organized within trypanosomatid PTUs, we analyzed three classes of functional relationships: subunits of the same protein complex, enzymes of the same metabolic pathway, and genes assigned to the same broad KEGG functional category.

Protein-complex annotations were obtained from the Complex Portal database [112]. The dataset included annotated subunit compositions for 2,420 human complexes, 752 mouse complexes, 631 yeast complexes, 269 fruit fly complexes, 225 *Arabidopsis thaliana* complexes, and 144 *Caenorhabditis elegans* complexes. Amino-acid sequences for complex subunits were retrieved from UniProt [161] and mapped to the protein families inferred in this study using reciprocal BLAST searches [21]. Hits were retained using an e-value cutoff of ≤10^−10^ (**Data S11**). In trypanosomatid genomes, tandem gene duplications frequently amplify gene dosage; these duplications can also drive false positive signatures of adjacency of functionally related genes. To account for this, for each complex subunit, only the best hit in the *L. major* genome (lowest e-value) was considered.

Metabolic-pathway annotations were obtained from MetaCyc [113]. For each annotated eukaryotic pathway, enzyme compositions were mapped to the protein-family annotations using the same reciprocal BLAST-based procedure described above. Additionally, the *L. major* proteome was used to identify genes related to broad functional classes. These classes were assigned using KEGG annotations [116] (**Data S12**).

For each protein complex, metabolic pathway, or KEGG functional class, we enumerated all pairs of *L. major* genes assigned to the same functional unit. Gene pairs were then classified according to whether both genes resided within the same annotated PTU. To test whether functionally linked gene pairs were enriched within shared PTUs, we compared the observed fraction of same-PTU pairs to a random background generated by sampling 10,000 random gene pairs from all annotated *L. major* genes. Enrichment was quantified as a Z-score relative to this random background.

For functionally linked gene pairs that resided within the same PTU, we next quantified their relative distance along the PTU. Gene-index distance was calculated as the absolute difference between the two genes’ positions within the PTU and was normalized by the total number of genes in that PTU. This normalization enabled comparison across PTUs of different lengths. The observed distribution of normalized gene-index distances for co-complex, co-pathway, or same-KEGG-class gene pairs was compared to a random background of within-PTU gene pairs. Random within-PTU pairs were sampled with probability weighted by PTU size, so that longer PTUs contributed more candidate pairs proportionally. Observed and random distance distributions were compared using two-sided Mann-Whitney U tests. Smaller normalized gene-index distances indicate greater adjacency or near-adjacency within shared PTUs.

