## Supplementary Information for "Evolution of a genome-architecture-encoded gene regulation system in trypanosomatids"

**Author Affiliations**

* Corresponding authors

**This Supplementary Materials file includes**

- Supplementary Data File Legends (S1-S12)
- Supplementary Tables S1-S4
- Supplementary Figures, S1-S4
- Supplementary References 1-65

**Supplementary Data File Legends**

**Data S1.** FASTA-format predicted proteomes for the 59 protist species analyzed in this study.

**Data S2.** Protein-family assignments inferred across the 59 protist proteomes.

**Data S3.** FASTA-format amino acid sequences of the 154 conserved, nearly single-copy protein families used for species-tree reconstruction.

**Data S4.** Average direction lengths for 11,477 genomes sampled across the Tree of Life.

**Data S5.** Genome annotations, in GFF3 format, for the 47 protist species with available genome assemblies. These annotations were used to identify directons and to extract inter- and intra-directon non-coding regions.

**Data S6.** Transcription-factor binding motifs identified as significantly enriched in inter-directon non-coding regions of the Discoba genomes analyzed in this study.

**Data S7.** Polycistronic transcription unit (PTU) definitions for the *Leishmania major* genome used in this study.

**Data S8.** Per-gene relative RNA polymerase II occupancy values for *Trypanosoma brucei*.

**Data S9.** Polycistronic transcription unit (PTU) definitions for the *Trypanosoma brucei* genome used in this study.

**Data S10.** Per-gene tRNA adaptation index (tAI) values calculated for the *Leishmania major* and *Trypanosoma brucei* genomes.

**Data S11.** *Leishmania major* orthologs of genes encoding subunits of annotated eukaryotic multiprotein complexes and enzymes of annotated metabolic pathways.

**Data S12.** *Leishmania major* genes assigned to broad functional categories based on KEGG annotations.

**Supplementary Tables**

**Table S1.** Taxon sampling, data sources, genome availability, and proteome-completeness metrics for the 59 protist species included in the phylogenetic analysis. For each species, the table lists the source publication or database, whether genome-assembly data were available, and BUSCO completeness scores.

| **Proteome** | **Reference** | **Data Type** | **BUSCO score** |
| --- | --- | --- | --- |
| *Acrasis kona* | Sheikh et al., 2024 | Genome | 85.5 |
| *Willaertia magna* | Zavadska et al. 2025 | Predicted Proteome | 85.1 |
| *Naegleria lovaniensis* | NCBI Genome | Genome | 86.3 |
| *Naegleria gruberi* | EukProt; NCBI Genome | Genome | 77.6 |
| *Neovahlkampfia damariscottae* | EukProt; NCBI Genome | Genome | 86.7 |
| *Pharyngomonas kirbyi* | EukProt; Benz et al. 2024 | Predicted Proteome | 84.7 |
| *Stygiella incarcerata* | EukProt; Benz et al. 2024 | Predicted Proteome | 69.0 |
| *Ophirina chinija* | Zavadska et al. 2025 | Predicted Proteome | 80.8 |
| *Ploeotia vitrea* | Benz et al. 2024 | Predicted Proteome | 58.8 |
| *Entosiphon sulcatum* | EukProt; Benz et al. 2024 | Predicted Proteome | 68.2 |
| *Rhabdomonas costata* | Zavadska et al. 2025 | Genome | 61.6 |
| *Euglena longa* | EukProt; Zavadska et al. 2025 | Predicted Proteome | 83.5 |
| *Euglena gracilis* | EukProt; Zavadska et al. 2025 | Genome | 71.0 |
| *Hemistasia phaeocysticola* | EukProt; Benz et al. 2024 | Predicted Proteome | 68.2 |
| *Sulcionema specki* | EukProt; Benz et al. 2024 | Predicted Proteome | 61.6 |
| *Rhynchopus humris* | EukProt; Zavadska et al. 2025 | Genome | 68.6 |
| *Diplonema papillatum* | NCBI Genome | Genome | 69.0 |
| *Perkinsela sp* | TryTripDB database | Genome | 65.4 |
| *Papus ankaliazontas* | Zavadska et al. 2025 | Predicted Proteome | 65.4 |
| *Apiculatamorpha spiralis* | Benz et al. 2024 | Predicted Proteome | 63.8 |
| *Neobodo designis* | EukProt; Zavadska et al. 2025 | Predicted Proteome | 51.5 |
| *Bodo saltans* | EukProt; Kostygov et al. 2024 | Genome | 87.7 |
| *Paratrypanosoma confusum* | Kostygov et al. 2024 | Genome | 96.2 |
| *Trypanosoma theileri* | VEuPathDB; Kostygov et al. 2024 | Genome | 99.2 |
| *Trypanosoma cruzi* | VEuPathDB; Kostygov et al. 2024 | Genome | 100.0 |
| *Trypanosoma vivax* | VEuPathDB; Kostygov et al. 2024 | Genome | 94.6 |
| *Trypanosoma brucei* | VEuPathDB; Kostygov et al. 2024 | Genome | 99.2 |
| *Blechomonas ayalai* | VEuPathDB; Kostygov et al. 2024 | Genome | 99.2 |
| *Vickermania ingenoplastis* | Kostygov et al. 2024 | Genome | 80.8 |
| *Obscuromonas volfi* | Kostygov et al. 2024 | Genome | 86.2 |
| *Obscuromonas eliasi* | Kostygov et al. 2024 | Genome | 88.5 |
| *Phytomonas sp* | Kostygov et al. 2024 | Genome | 90.0 |
| *Phytomonas francai* | Kostygov et al. 2024 | Genome | 73.1 |
| *Herpetomonas tarakana* | Kostygov et al. 2024 | Genome | 95.4 |
| *Herpetomonas samuelpessoai* | Kostygov et al. 2024 | Genome | 99.2 |
| *Wallacemonas collosoma* | Kostygov et al. 2024 | Genome | 98.5 |
| *Wallacemonas rigidus* | Kostygov et al. 2024 | Genome | 98.5 |
| *Kentomonas sorsogonicus* | Kostygov et al. 2024 | Genome | 70.8 |
| *Strigomonas oncopelti* | Kostygov et al. 2024 | Genome | 91.5 |
| *Strigomonas galati* | Kostygov et al. 2024 | Genome | 90.0 |
| *Angomonas ambiguus* | Kostygov et al. 2024 | Genome | 76.9 |
| *Angomonas deanei* | VEuPathDB; Kostygov et al. 2024 | Genome | 77.7 |
| *Sergeia podlipaevii* | Kostygov et al. 2024 | Genome | 90.0 |
| *Jaenimonas drosophilae* | Kostygov et al. 2024 | Genome | 91.5 |
| *Crithidia thermophila* | Kostygov et al. 2024 | Genome | 96.9 |
| *Leptomonas seymouri* | VEuPathDB; Kostygov et al. 2024 | Genome | 99.2 |
| *Leptomonas pyrrhocoris* | VEuPathDB; Kostygov et al. 2024 | Genome | 100.0 |
| *Lotmaria passim* | Kostygov et al. 2024 | Genome | 100.0 |
| *Crithidia expoeki* | Kostygov et al. 2024 | Genome | 96.2 |
| *Borovskyia barvae* | Kostygov et al. 2024 | Genome | 97.7 |
| *Zelonia costaricensis* | Kostygov et al. 2024 | Genome | 96.9 |
| *Novymonas esmeraldas* | Kostygov et al. 2024 | Genome | 96.2 |
| *Endotrypanum monterogeii* | VEuPathDB; Kostygov et al. 2024 | Genome | 97.7 |
| *Porcisia hertigi* | VEuPathDB; Kostygov et al. 2024 | Genome | 98.5 |
| *Porcisia deanei* | Kostygov et al. 2024 | Genome | 98.5 |
| *Leishmania martiniquensis* | VEuPathDB; Kostygov et al. 2024 | Genome | 100.0 |
| *Leishmania panamensis* | VEuPathDB; Kostygov et al. 2024 | Genome | 98.5 |
| *Leishmania major* | VEuPathDB; Kostygov et al. 2024 | Genome | 100.0 |
| *Leishmania donovani* | VEuPathDB; Kostygov et al. 2024 | Genome | 100.0 |

**Table S2.** Pfam-annotated DNA-binding domains associated with transcription-factor activity. These domains were used to identify proteins harboring transcription-factor-associated DNA-binding domains across the 59 protist proteomes.

| **Pfam Accession** | **Short Name** | **Long Name** |
| --- | --- | --- |
| PF00010 | HLH | Helix-loop-helix DNA-binding domain |
| PF00216 | Bac_DNA_binding | Bacterial DNA-binding protein |
| PF00249 | Myb_DNA-binding | Myb-like DNA-binding domain |
| PF00319 | SRF-TF | SRF-type transcription factor (DNA-binding and dimerisation domain) |
| PF00447 | HSF_DNA-bind | HSF-type DNA-binding |
| PF00554 | RHD_DNA_bind | Rel homology DNA-binding domain |
| PF00649 | Copper-fist | Copper fist DNA binding domain |
| PF00751 | DM | DM DNA binding domain |
| PF00870 | P53 | P53 DNA-binding domain |
| PF01316 | Arg_repressor | Arginine repressor, DNA binding domain |
| PF01325 | Fe_dep_repress | Iron dependent repressor, N-terminal DNA binding domain |
| PF01388 | ARID | ARID/BRIGHT DNA binding domain |
| PF01726 | LexA_DNA_bind | LexA DNA binding domain |
| PF02257 | RFX_DNA_binding | RFX DNA-binding domain |
| PF02319 | WHD_E2F_TDP | E2F/DP family winged-helix DNA-binding domain |
| PF02362 | B3 | B3 DNA binding domain |
| PF02864 | STAT_bind | STAT protein, DNA binding domain |
| PF03101 | FAR1 | FAR1 DNA-binding domain |
| PF03106 | WRKY | WRKY DNA -binding domain |
| PF03444 | WHD_HrcA | Winged helix-turn-helix transcription repressor, HrcA DNA-binding |
| PF04552 | Sigma54_DBD | Sigma-54, DNA binding domain |
| PF04873 | EIN3_DNA-bd | Ethylene insensitive 3, DNA-binding domain |
| PF04967 | HTH_10 | HTH DNA binding domain |
| PF05224 | NDT80_PhoG | NDT80 / PhoG like DNA-binding family |
| PF07842 | GCFC | GC-rich sequence DNA-binding factor-like protein |
| PF07879 | PHB_acc_N | PHB/PHA accumulation regulator DNA-binding domain |
| PF09270 | BTD | Beta-trefoil DNA-binding domain |
| PF09278 | MerR-DNA-bind | MerR, DNA binding |
| PF09287 | CEP1-DNA_bind | CEP-1, DNA binding |
| PF09607 | BrkDBD | Brinker DNA-binding domain |
| PF10074 | RovC_DNA-bd | T6SS, Transcription factor, DNA binding domain |
| PF10416 | IBD | Transcription-initiator DNA-binding domain IBD |
| PF10491 | Nrf1_DNA-bind | NLS-binding and DNA-binding and dimerisation domains of Nrf1 |
| PF10524 | NfI_DNAbd_pre-N | Nuclear factor I protein pre-N-terminus |
| PF10545 | MADF_DNA_bdg | Alcohol dehydrogenase transcription factor Myb/SANT-like |
| PF11972 | HTH_13 | HTH DNA binding domain |
| PF12181 | MogR_DNAbind | DNA binding domain of the motility gene repressor (MogR) |
| PF12329 | TMF_DNA_bd | TATA element modulatory factor 1 DNA binding |
| PF12776 | Myb_DNA-bind_3 | Myb/SANT-like DNA-binding domain |
| PF13412 | HTH_24 | Winged helix-turn-helix DNA-binding |
| PF13443 | HTH_26 | Cro/C1-type HTH DNA-binding domain |
| PF13463 | HTH_27 | Winged helix DNA-binding domain |
| PF13601 | HTH_34 | Winged helix DNA-binding domain |
| PF13693 | HTH_35 | Winged helix-turn-helix DNA-binding |
| PF13837 | Myb_DNA-bind_4 | Myb/SANT-like DNA-binding domain |
| PF13873 | Myb_DNA-bind_5 | Myb/SANT-like DNA-binding domain |
| PF13921 | Myb_DNA-bind_6 | Myb-like DNA-binding domain |
| PF14621 | RFX5_DNA_bdg | RFX5 DNA-binding domain |
| PF14850 | Pro_dh-DNA_bdg | DNA-binding domain of Proline dehydrogenase |
| PF15963 | Myb_DNA-bind_7 | Myb DNA-binding like |
| PF15977 | HTH_46 | Winged helix-turn-helix DNA binding |
| PF16422 | COE1_DBD | Transcription factor COE1 DNA-binding domain |
| PF16557 | CUTL | CUT1-like DNA-binding domain of SATB |
| PF16721 | zf-H3C2 | Zinc-finger like, probable DNA-binding |
| PF17538 | C_LFY_FLO | DNA Binding Domain (C-terminal) Leafy/Floricaula |
| PF21179 | BldD-like_C | DNA-binding protein BldD-like, C-terminal domain |
| PF21227 | Myb_DNA-binding_7 | Myb-like DNA-binding domain |
| PF21321 | HTH_66 | Putative DNA-binding HTH domain |
| PF21411 | Rap1_C_Sp | S. pombe DNA-binding protein Rap1, C-terminal |
| PF21559 | Reb1_MybAD | DNA-binding protein reb1, Myb-associated domain |
| PF21715 | CggR_N | CggR N-terminal DNA binding domain |
| PF21906 | WHD_NrtR | NrtR DNA-binding winged helix domain |
| PF22980 | Myb_DNA-bind_8 | Myb-like DNA-binding domain |
| PF23082 | Myb_DNA-binding_2 | Myb-like DNA-binding domain |
| PF25603 | SPT23_MGA2_DBD | SPT23/MGA2 DNA-binding domain |

**Table S3.** Pfam-annotated RNA-binding domains associated with post-transcriptional regulation, including mRNA stability, localization, and translational control. These domains were used to identify proteins harboring RNA-binding domains across the 59 protist proteomes.

| **Pfam Accession** | **Short Name** | **Long Name** |
| --- | --- | --- |
| PF00035 | dsrm | Double-stranded RNA binding motif |
| PF00074 | RnaseA | Pancreatic ribonuclease |
| PF00075 | RNase_H | RNase H |
| PF00076 | RRM_1 | RNA recognition motif |
| PF00806 | PUF | Pumilio-family RNA binding repeat |
| PF00910 | RNA_helicase | RNA helicase |
| PF00929 | RNase_T | Exonuclease |
| PF01125 | BUD31 | Pre-mRNA-splicing factor BUD31 |
| PF01138 | RNase_PH | 3' exoribonuclease family, domain 1 |
| PF01223 | Endonuclease_NS | DNA/RNA non-specific endonuclease |
| PF01351 | RNase_HII | Ribonuclease HII |
| PF01868 | RNase_P-MRP_p29 | Ribonuclease P/MRP, subunit p29 |
| PF01876 | RNase_P_p30 | RNase P subunit p30 |
| PF01877 | RNA_binding | RNA binding |
| PF01900 | RNase_P_Rpp14 | Rpp14/Pop5 family |
| PF01927 | Mut7-C | Mut7-C RNAse domain |
| PF02081 | TrpBP | Tryptophan RNA-binding attenuator protein |
| PF03123 | CAT_RBD | CAT RNA binding domain |
| PF03725 | RNase_PH_C | 3' exoribonuclease family, domain 2 |
| PF03726 | PNPase | Polyribonucleotide nucleotidyltransferase, RNA binding domain |
| PF04032 | Rpr2 | RNAse P Rpr2/Rpp21/SNM1 subunit domain |
| PF04059 | RRM_2 | RNA recognition motif 2 |
| PF04135 | Nop10p | Nucleolar RNA-binding protein, Nop10p family |
| PF04308 | RNaseH_like | Ribonuclease H-like |
| PF04410 | Gar1 | Gar1/Naf1 RNA binding region |
| PF04774 | HABP4_PAI-RBP1 | Hyaluronan / mRNA binding family |
| PF04926 | PAP_RNA-bind | Poly(A) polymerase predicted RNA binding domain |
| PF05172 | RRM_Nup35 | Nup53/35/40-type RNA recognition motif |
| PF05634 | APO_RNA-bind | APO RNA-binding |
| PF05652 | DcpS | Scavenger mRNA decapping enzyme (DcpS) N-terminal |
| PF05670 | NFACT-R_1 | NFACT protein RNA binding domain |
| PF05741 | zf-nanos | Nanos RNA binding domain |
| PF06485 | Tab2-like_N | RNA-binding protein Tab2/Atab2 N terminal |
| PF06807 | Clp1 | Pre-mRNA cleavage complex II protein Clp1 |
| PF06991 | MFAP1 | Microfibril-associated/Pre-mRNA processing |
| PF07497 | Rho_RNA_bind | Rho termination factor, RNA-binding domain |
| PF08228 | RNase_P_pop3 | RNase P subunit Pop3 |
| PF08315 | cwf18 | cwf18 pre-mRNA splicing factor |
| PF08424 | NRDE-2 | NRDE-2, necessary for RNA interference |
| PF08459 | UvrC_RNaseH_dom | UvrC RNAse H endonuclease domain |
| PF08461 | WHD_RNase_R | Ribonuclease R winged-helix domain |
| PF08572 | PRP3 | pre-mRNA processing factor 3 domain |
| PF08615 | RNase_H2_suC | Ribonuclease H2 non-catalytic subunit (Ylr154p-like) |
| PF08675 | RNA_bind | RNA binding domain |
| PF08777 | RRM_3 | RNA binding motif |
| PF08799 | PRP4 | pre-mRNA processing factor 4 (PRP4) like |
| PF09162 | Tap-RNA_bind | Tap, RNA-binding |
| PF09387 | MRP | Mitochondrial RNA binding protein MRP |
| PF09416 | UPF1_Zn_bind | RNA helicase (UPF2 interacting domain) |
| PF09468 | RNase_H2-Ydr279 | Ydr279p protein family (RNase H2 complex component) wHTH domain |
| PF09692 | Arb1 | Argonaute siRNA chaperone (ARC) complex subunit Arb1 |
| PF09736 | Bud13 | Pre-mRNA-splicing factor of RES complex |
| PF09750 | DRY_EERY | Alternative splicing regulator |
| PF09848 | SLFN-g3_helicase | Schlafen group 3, DNA/RNA helicase domain |
| PF10150 | RNase_E_G | Ribonuclease E/G family |
| PF10258 | PHAX_RNA-bd | Phosphorylated adapter RNA export protein, RNA-binding domain |
| PF10429 | Mtr2 | Nuclear pore RNA shuttling protein Mtr2 |
| PF10443 | RNA12 | RNA12 protein |
| PF10500 | SR-25 | Nuclear RNA-splicing-associated protein |
| PF10567 | Nab6_mRNP_bdg | RNA-recognition motif |
| PF10598 | RRM_4 | RNA recognition motif of the spliceosomal PrP8 |
| PF11435 | She2p | RNA binding protein She2p |
| PF11473 | B2 | RNA binding protein B2 |
| PF11517 | Nab2 | Nuclear abundant poly(A) RNA-bind protein 2 (Nab2) |
| PF11608 | RRM_MARF1 | MARF1, RNA recognition motif 1 |
| PF11708 | Slu7 | Pre-mRNA splicing Prp18-interacting factor |
| PF11717 | Tudor-knot | RNA binding activity-knot of a chromodomain |
| PF11718 | CPSF73-100_C | Pre-mRNA 3'-end-processing endonuclease polyadenylation factor C-term |
| PF11831 | Myb_Cef | pre-mRNA splicing factor component |
| PF11969 | DcpS_C | Scavenger mRNA decapping enzyme C-term binding |
| PF11977 | RNase_Zc3h12a | Zc3h12a-like Ribonuclease NYN domain |
| PF12011 | NPH-II | RNA helicase NPH-II |
| PF12072 | RNase_Y_N | RNase Y N-terminal region |
| PF12171 | zf-C2H2_jaz | Zinc-finger double-stranded RNA-binding |
| PF12230 | PRP21_like_P | Pre-mRNA splicing factor PRP21 like protein |
| PF12300 | RhlB | ATP-dependent RNA helicase RhlB |
| PF12328 | Rpp20 | Rpp20 subunit of nuclear RNase MRP and P |
| PF12513 | SUV3_C | ATP-dependent RNA helicase SUV3 C-terminal domain |
| PF12542 | CWC25 | Pre-mRNA splicing factor |
| PF12622 | NpwBP | mRNA biogenesis factor |
| PF12871 | PRP38_assoc | Pre-mRNA-splicing factor 38-associated hydrophilic C-term |
| PF13017 | Maelstrom | piRNA pathway germ-plasm component |
| PF13032 | RNaseH_pPIWI_RE | RNaseH domain of pPIWI_RE |
| PF13482 | RNase_H_2 | RNase_H superfamily |
| PF13893 | RRM_5 | RNA recognition motif. |
| PF13929 | mRNA_stabil | mRNA stabilisation |
| PF13965 | SID-1_RNA_chan | dsRNA-gated channel SID-1 |
| PF14605 | Nup35_RRM_2 | Nup53/35/40-type RNA recognition motif |
| PF14608 | zf-CCCH_2 | RNA-binding, Nab2-type zinc finger |
| PF14626 | RNase_Zc3h12a_2 | Zc3h12a-like Ribonuclease NYN domain |
| PF14709 | DND1_DSRM | double strand RNA binding domain from DEAD END PROTEIN 1 |
| PF15247 | SLBP_RNA_bind | Histone RNA hairpin-binding protein RNA-binding domain |
| PF15608 | PELOTA_1 | PELOTA RNA binding domain |
| PF15777 | Anti-TRAP | Tryptophan RNA-binding attenuator protein inhibitory protein |
| PF15801 | zf-C6H2 | zf-MYND-like zinc finger, mRNA-binding |
| PF16005 | MOEP19 | KH-like RNA-binding domain |
| PF16367 | RRM_7 | RNA recognition motif |
| PF16529 | Ge1_WD40 | WD40 region of Ge1, enhancer of mRNA-decapping protein |
| PF16575 | CLP1_P | mRNA cleavage and polyadenylation factor CLP1 P-loop |
| PF16598 | Edc3_linker | Linker region of enhancer of mRNA-decapping protein 3 |
| PF16741 | mRNA_decap_C | mRNA-decapping enzyme C-terminus |
| PF16835 | SF3A2 | Pre-mRNA-splicing factor SF3a complex subunit 2 (Prp11) |
| PF16837 | SF3A3 | Pre-mRNA-splicing factor SF3A3, of SF3a complex, Prp9 |
| PF16842 | RRM_occluded | Occluded RNA-recognition motif |
| PF16953 | PRORP | Protein-only RNase P |
| PF16958 | PRP9_N | Pre-mRNA-splicing factor PRP9 N-terminus |
| PF16969 | SRP68 | RNA-binding signal recognition particle 68 |
| PF17208 | RBR | RNA binding Region |
| PF17331 | GFD1 | GFD1 mRNA transport factor |
| PF17587 | Dmd | Discriminator of mRNA degradation |
| PF17703 | C19orf84 | piRNA-mediated silencing protein C19orf84 |
| PF17770 | RNase_J_C | Ribonuclease J C-terminal domain |
| PF17774 | YlmH_RBD | Putative RNA-binding domain in YlmH |
| PF17842 | dsRBD2 | Double-stranded RNA binding domain 2 |
| PF18141 | UPF1_1B_dom | RNA helicase UPF1, 1B domain |
| PF18260 | Nab2p_Zf1 | Nuclear polyadenylated RNA-binding 2 protein CCCH zinc finger 1 |
| PF18297 | NFACT-R_2 | NFACT protein RNA binding domain |
| PF18431 | RNAse_A_bac | Bacterial CdiA-CT RNAse A domain |
| PF18444 | RRM_9 | RNA recognition motif |
| PF18497 | RNase_3_N | Ribonuclease III N-terminal domain |
| PF18528 | Ret2_MD | RNA editing 3' terminal uridylyl transferase 2 middle domain |
| PF18614 | RNase_II_C_S1 | RNase II-type exonuclease C-terminal S1 domain |
| PF18869 | HEPN_RnaseLS | RnaseLS-like HEPN |
| PF19034 | RnlA-toxin_DBD | RNase LS, bacterial toxin DBD domain |
| PF19417 | RnlA_toxin_N | RNase LS, bacterial toxin N-terminal |
| PF20429 | Tab2-like_C | RNA-binding protein Tab2/Atab2, C-terminal domain |
| PF20822 | Cmr2_hel_dom1 | CRISPR RNA silencing complex Cmr2 subunit, first helical domain |
| PF20823 | Cmr2_Zn-bd | CRISPR RNA silencing complex Cmr2 subunit, Zn-binding domain |
| PF20824 | Cmr2_hel_dom2 | CRISPR RNA silencing complex Cmr2 subunit, second helical domain |
| PF20833 | RNase_E_G_Thio | RNase E/G, Thioredoxin-like domain |
| PF20844 | PCF11_RFEG_rpt | Pre-mRNA cleavage complex 2 protein Pcf11, RFEGP repeat |
| PF20845 | Pcf11_helical | Pre-mRNA cleavage complex 2 protein Pcf11, alpha-helical domain |
| PF20932 | Dicer_dsRBD | Dicer, dsRNA-binding domain |
| PF21056 | ZSWIM1-3_RNaseH-like | Zinc finger SWIM domain-containing protein 1/3, RNaseH-like domain |
| PF21188 | BRR2_plug | Pre-mRNA-splicing helicase BRR2 plug domain |
| PF21210 | RNA_helicase_helical | Putative ATP-dependent RNA helicase, helical bundle |
| PF21289 | EDC4_C | Enhancer of mRNA-decapping protein 4, C-terminal |
| PF21293 | RNAseD_HRDC_C | Ribonuclease D, C-terminal HRDC domain |
| PF21408 | MTR4-like_stalk | Exosome RNA helicase MTR4-like, stalk |
| PF21457 | zf-CCCH_2-like_3 | RNA-binding, Nab2-type zinc finger |
| PF21591 | Hera_DD | RNA helicase Hera, dimerization domain |
| PF21787 | TNP-like_RNaseH_N | TNP-like, RNase H N-terminal domain |
| PF21789 | TNP-like_RNaseH_C | TNP-like, RNase H C-terminal domain |
| PF21864 | MORF_dom | Multiple organellar RNA editing factor 1, MORF domain |
| PF21940 | Pfc11_Rna14-15-ID | Pfc11 Rna14/15 interacting domain |
| PF22231 | Hera_RBD | DEAD box helicase Hera, RNA-binding domain |
| PF22327 | Nudt16-like | U8 snoRNA-decapping enzyme-like |
| PF22389 | Dcr1-like_dsRNA-bd_dom | Protein Dicer, dsRNA-binding domain |
| PF22505 | RNase_J_b_CASP | Ribonuclease J, beta-CASP domain |
| PF22602 | NXF_NTF2 | Nuclear RNA export factor, NTF2 domain |
| PF22625 | ECR1_N_2 | Exosome RNA binding protein RRP4 N-terminal domain |
| PF22892 | DSRM_MRPL44 | MRPL44 dsRNA-binding domain |
| PF22901 | dsrm_Ferlin | Ferlin dsRNA-binding domain-like domain |
| PF22920 | UvrC_RNaseH | UvrC Ribonuclease H-like domain |
| PF23002 | PIN-like_DDX60 | ATP-dependent RNA helicase DDX60, PIN-like domain |
| PF23057 | RBD_ZCCHC3_1st | ZCCHC3, first RNA-binding domain |
| PF23058 | RBD_ZCCHC3_2nd | ZCCHC3, second RNA-binding domain |
| PF23161 | HTH_RNase_II | Double HTH domain |
| PF23163 | CSD_RNase_II | CSD-like barrel domain |
| PF23220 | HAT_Syf1_M | Pre-mRNA-splicing factor SYF1 middle HAT repeat |
| PF23231 | HAT_Syf1_CNRKL1_C | Pre-mRNA-splicing factor Syf1/CRNKL1 C-terminal HAT repeat |
| PF23233 | HAT_Syf1_CNRKL1_N | Pre-mRNA-splicing factor Syf1/CNRKL1 N-terminal HAT repeat |
| PF23703 | DExH18_N | DExH-box ATP-dependent RNA helicase DExH18, N-terminal |
| PF24099 | RBD_DGKtheta | Diacylglycerol kinase theta, RNA-binding domain-like |
| PF24385 | DSRM_DHX29 | ATP-dependent RNA helicase DHX29, DSRM-like |
| PF24475 | RBD_DEAH11 | DEAH11-like, RNA-binding domain |
| PF24572 | RBD_RDR6 | RNA-dependent RNA polymerase 6, RNA-binding domain |
| PF24825 | SNU71_RBD | SNU71 RNA binding domain |
| PF24899 | UBA_DHX29 | ATP-dependent RNA helicase DHX29, UBA domain |
| PF24995 | DSRM_2 | dsRNA binding domain |
| PF25015 | RBD_AKAP-17A | AKAP-17A-like, RNA-binding domain |
| PF25061 | RRM_fung | Fungal RNA-binding domain |
| PF25255 | WHD_RNase_II | Ribonuclease II, winged helix domain |
| PF25399 | DeaD_dimer | RNA helicase DeaD dimerization domain |
| PF25479 | Vts1 | RNA-binding protein vts1-like domain |
| PF25488 | RNaseT2L_C | RNase T2-like C-terminal domain |
| PF26026 | RNA_hel_CTD | C-terminal domain in RNA helicases |
| PF26088 | RRM_LARP4 | La-related protein 4-like, RNA recognition motif |
| PF26165 | RESC7 | RNA-editing substrate-binding complex 7 protein |
| PF26170 | RESC5 | RNA-editing substrate-binding complex 5 protein |
| PF26172 | RESC8 | RNA-editing substrate-binding complex 8 protein |
| PF26179 | RESC10 | RNA-editing substrate-binding complex 10 protein |
| PF26188 | RESC6 | RNA-editing substrate-binding complex 6 protein |
| PF26980 | DSRM_DCL2 | Dicer-like protein 2, dsRNA binding domain |
| PF27148 | Slr1353_N | Slr1353 N-terminal dsRNA-binding domain-like |
| PF27457 | RNase_HII_N | Ribonuclease HII N-terminal domain |
| PF27771 | MP81 | RNA editing complex, structural subunit MP81 |
| PF28333 | RBM48_helical | RNA-binding protein 48 helical domain |
| PF28365 | REXO3_N | RNA exonuclease 3 N-terminal domain |
| PF28538 | Helicase_helical | Probable ATP-dependent RNA helicase, helical domain |

**Table S4.** Source organisms and references for the 10,607 RNA-seq datasets spanning 38 model organisms used in the cross-kingdom gene-activity analysis. Broad taxonomic groupings are provided for each organism.

| **Taxonomy** | **Species** | **Organism type** | **#datasets** | **References** |
| --- | --- | --- | --- | --- |
| Metazoa | *Homo sapiens* | Multicellular eukaryote | 1045 | Bgee |
|  | *Mus musculus* | Multicellular eukaryote | 1551 | Bgee |
|  | *Gallus gallus* | Multicellular eukaryote | 445 | Bgee |
|  | *Drosophila melanogaster* | Multicellular eukaryote | 555 | Bgee |
|  | *Caenorhabditis elegans* | Multicellular eukaryote | 149 | Bgee |
| Plants | *Zea mays* | Multicellular eukaryote | 1619 | Expression Atlas |
|  | *Oryza sativa* | Multicellular eukaryote | 444 | Expression Atlas |
|  | *Arabidopsis thaliana* | Multicellular eukaryote | 1470 | Expression Atlas |
|  | *Lycophytes* | Multicellular eukaryote | 35 | Ferrari et al., 2019 |
|  | *Mosses* | Multicellular eukaryote | 36 | Ferrari et al., 2019 |
| Algae | *Brown algae* | Multicellular eukaryote | 45 | Ferrari et al., 2019 |
|  | *Red algae* | Multicellular eukaryote | 91 | Ferrari et al., 2019 |
|  | *Green algae* | Multicellular eukaryote | 45 | Ferrari et al., 2019 |
| Fungi | *Saccharomyces cerevisiae* | Unicellular eukaryote | 1307 | Fungi.guru |
|  | *Schizosaccharomyces pombe* | Unicellular eukaryote | 557 | Fungi.guru |
|  | *Candida albicans* | Unicellular eukaryote | 62 | Fungi.guru |
|  | *Cryptococcus neoformans* | Unicellular eukaryote | 49 | Fungi.guru |
|  | *Sclerotinia sclerotiorum* | Multicellular eukaryote | 80 | Fungi.guru |
|  | *Puccinia striiformis* | Multicellular eukaryote | 49 | Fungi.guru |
| Protist | *Chlorella vulgaris* | Unicellular eukaryote | 30 | Protist.guru |
|  | *Chlamydomonas reinhardtii* | Unicellular eukaryote | 59 | Protist.guru |
|  | *Volvox carteri* | Unicellular eukaryote | 15 | Protist.guru |
| Trypanosomatids | *Leishmania major* | Unicellular eukaryote | 29 | Dillon et al., 2015;  Fernandes et al., 2016; Inbar et al., 2017 |
|  | *Trypanosoma brucei* | Unicellular eukaryote | 22 | Jensen et al., 2014;  Trindade et al., 2016 |
|  | *Trypanosoma cruzi* | Unicellular eukaryote | 24 | Inchausti et al. 2025 |
| Bacteria | *Escherichia coli* | Unicellular bacteria | 280 | Bacteria.guru |
|  | *Pseudomonas aeruginosa* | Unicellular bacteria | 28 | Bacteria.guru |
|  | *Vibrio cholerae* | Unicellular bacteria | 81 | Bacteria.guru |
|  | *Helicobacter pylori* | Unicellular bacteria | 29 | Bacteria.guru |
|  | *Campylobacter jejuni* | Unicellular bacteria | 48 | Bacteria.guru |
|  | *Neisseria gonorrhoeae* | Unicellular bacteria | 60 | Bacteria.guru |
|  | *Staphylococcus aureus* | Unicellular bacteria | 76 | Bacteria.guru |
|  | *Streptococcus pneumoniae* | Unicellular bacteria | 79 | Bacteria.guru |
|  | *Clostridioides difficile* | Unicellular bacteria | 60 | Bacteria.guru |
|  | *Mycobacterium tuberculosis* | Unicellular bacteria | 53 | Bacteria.guru |

**Supplementary Figures**

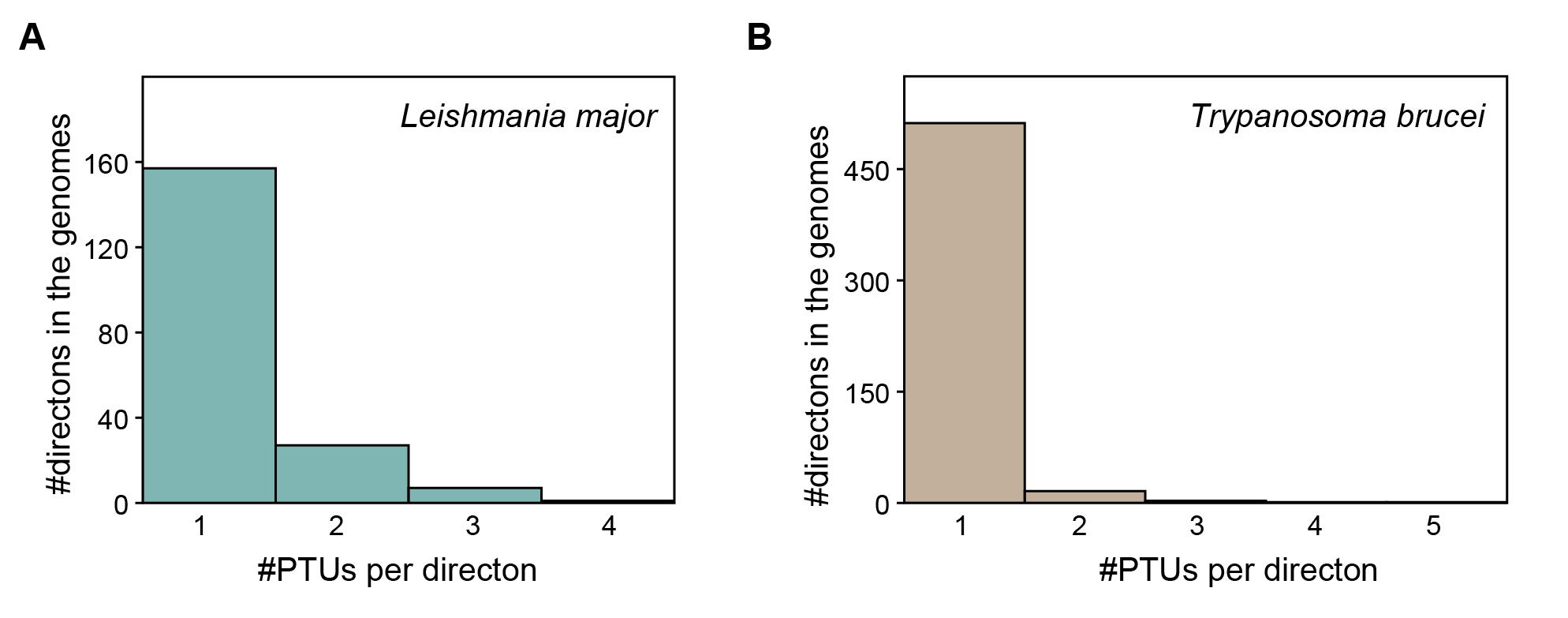

**Figure S1.** **Directons are a reliable proxy of trypanosomatids’ polycistronic gene arrangements.** (**A-B**) Histogram plots highlight the number of co-linear PTUs per directon for *L. major* (**A**) and *T. bruce*i genomes (**B**). These plots show that an overwhelming majority of directons on both genomes include a single PTU.

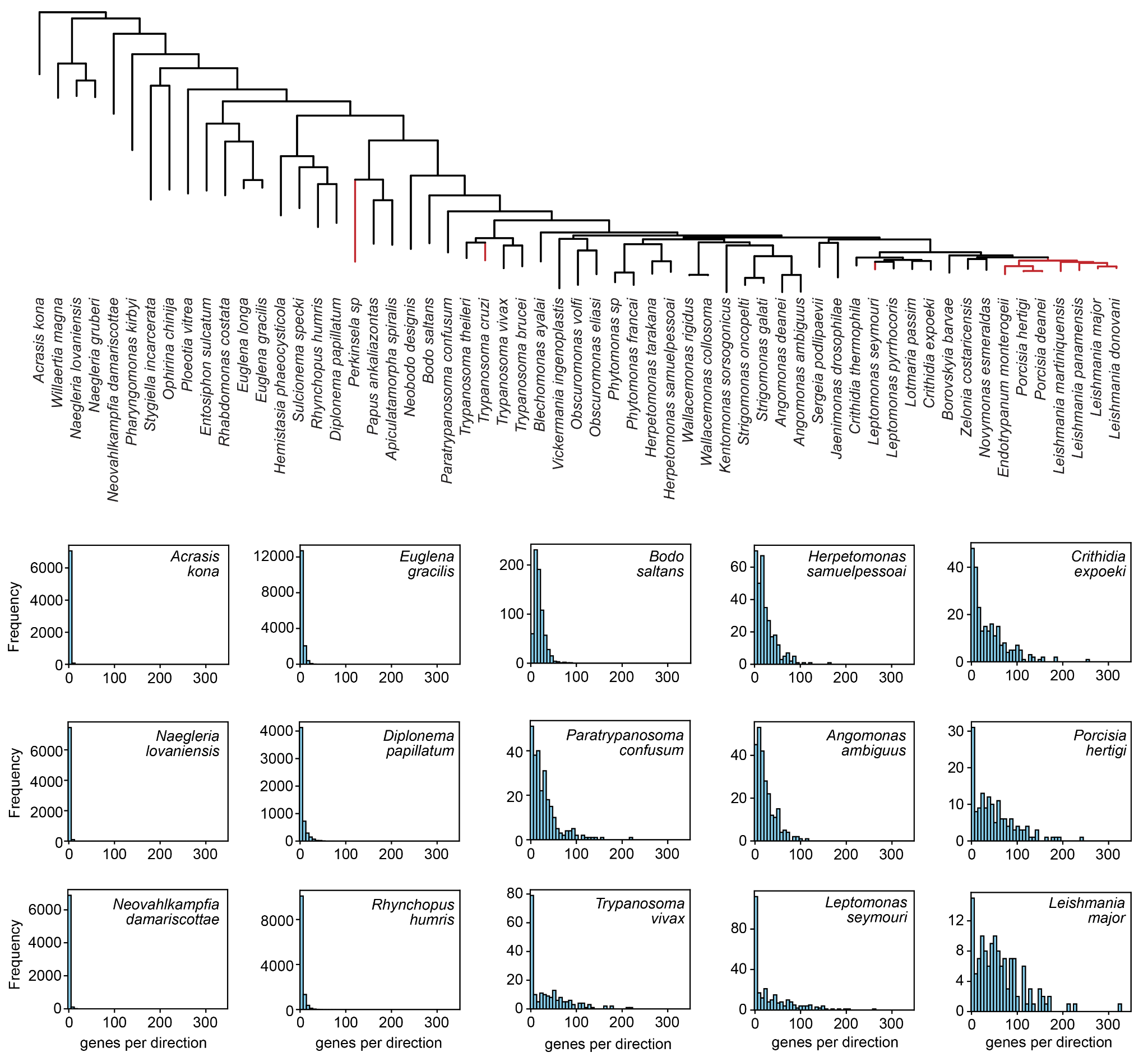

**Figure S2.** **Expansion of directional gene clusters in the Discoba lineage leading to *Leishmania*.** For representative genomes across the species tree, histogram plots represent the distribution of directon lengths (#genes per directon). The same X-axis range was used for each histogram. In the lineage leading to Leishmania we see longer directons progressively taking over the genomes.

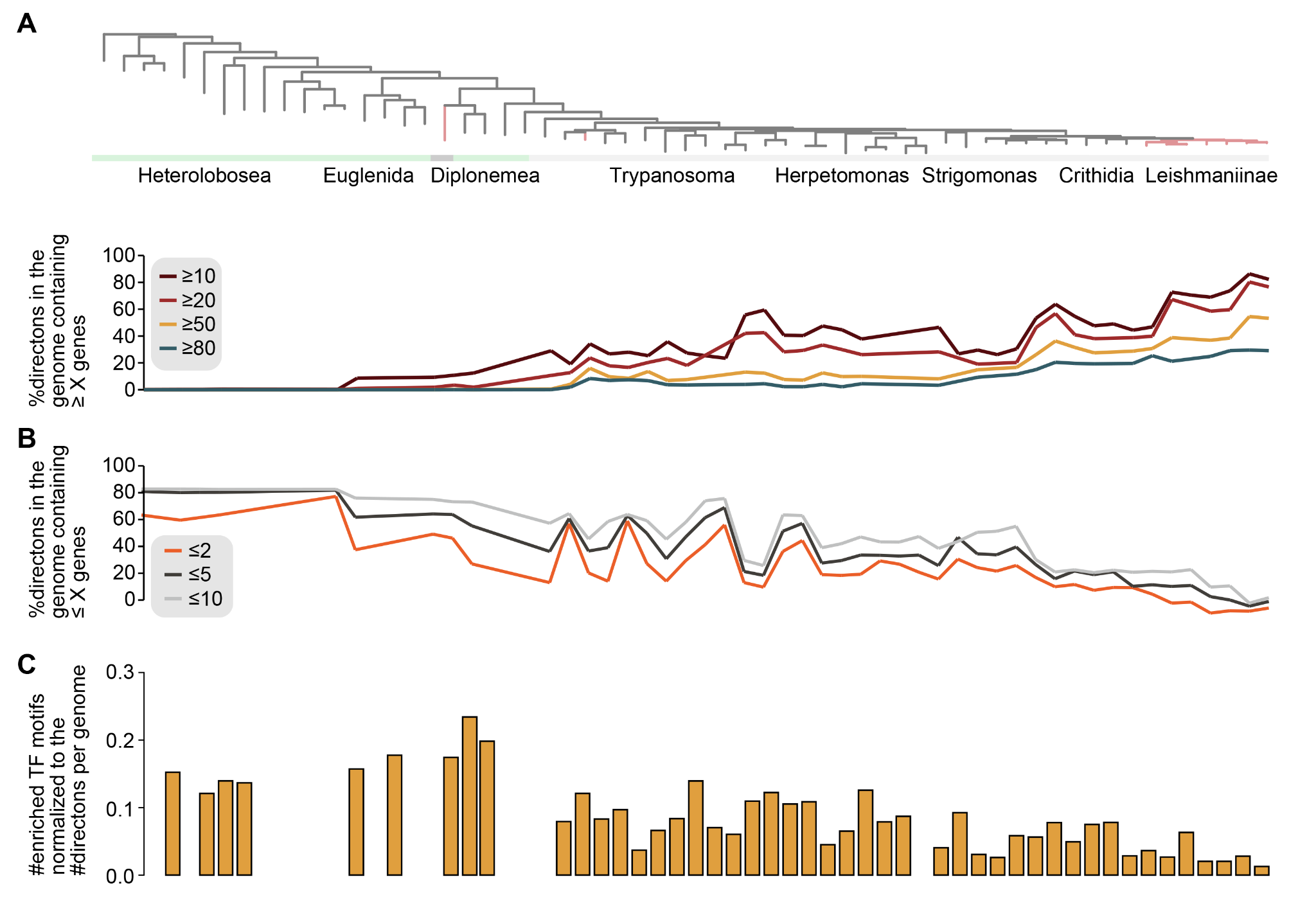

**Figure S3.** **Expansion of directional gene clusters and depletion of transcription-factor binding motifs across Discoba genomes.** (**A**) Lines show the percentage of directons containing at least X genes across the analyzed genomes. Different colors represent different X-value cutoffs: X ≥ 10, dark red; X ≥ 20, red; X ≥ 50, yellow; X ≥ 80, blue. (**B**) Lines show the percentage of directons containing at most Y genes. Different colors represent different Y-value cutoffs: Y ≤ 2, orange; Y ≤ 5, black; Y ≤ 10, gray. (**C**) Bar plots show the number of unique transcription-factor binding motifs (TFBMs) significantly enriched in inter-directon non-coding regions, normalized by the number of directons per genome.

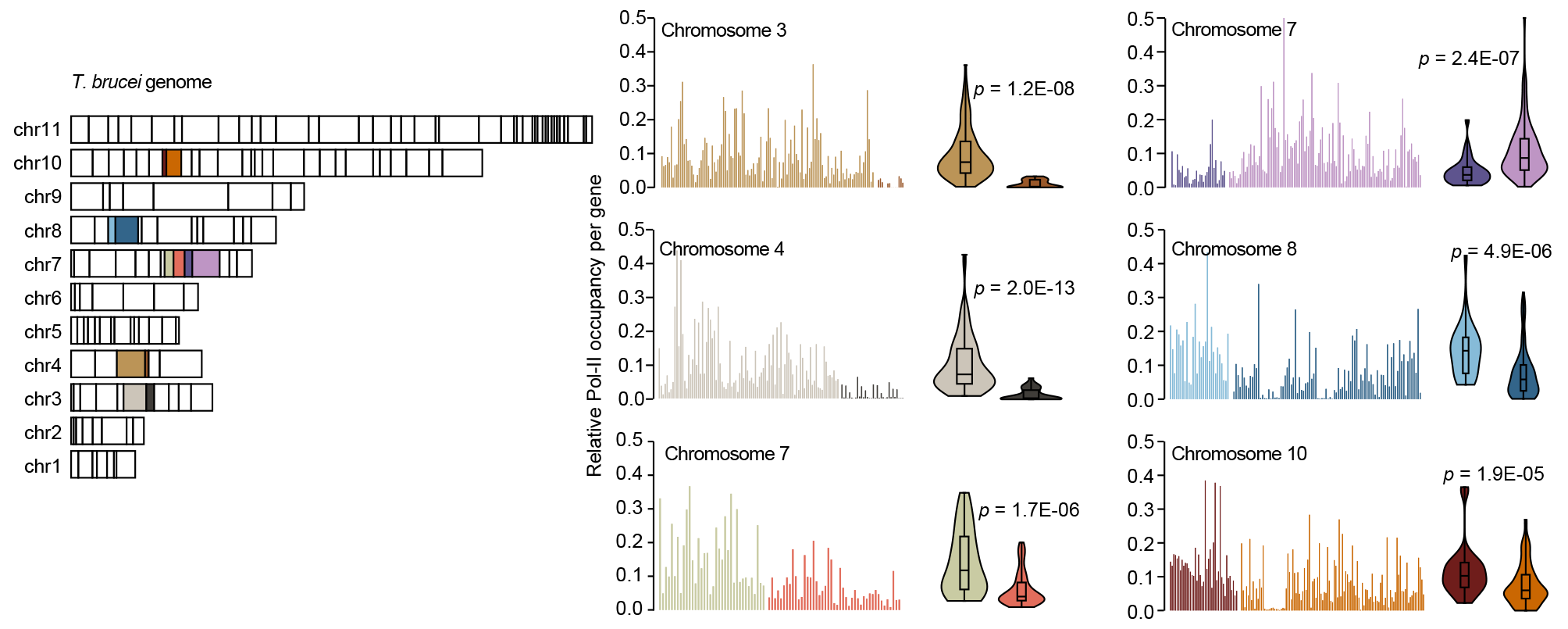
**Figure S4.** Same as **Figure 3F**, bar plots compare the per-gene relative per-gene Pol-II occupancy for neighboring PTU pairs in various chromosomes of *L. major* genome. The same data is also shown as violin plots, where each violin corresponds to a PTU. The *p*-values were derived from the Mann-Whitney U test. These plots demonstrate that neighboring PTUs often exhibit significantly different Pol-II occupancies.

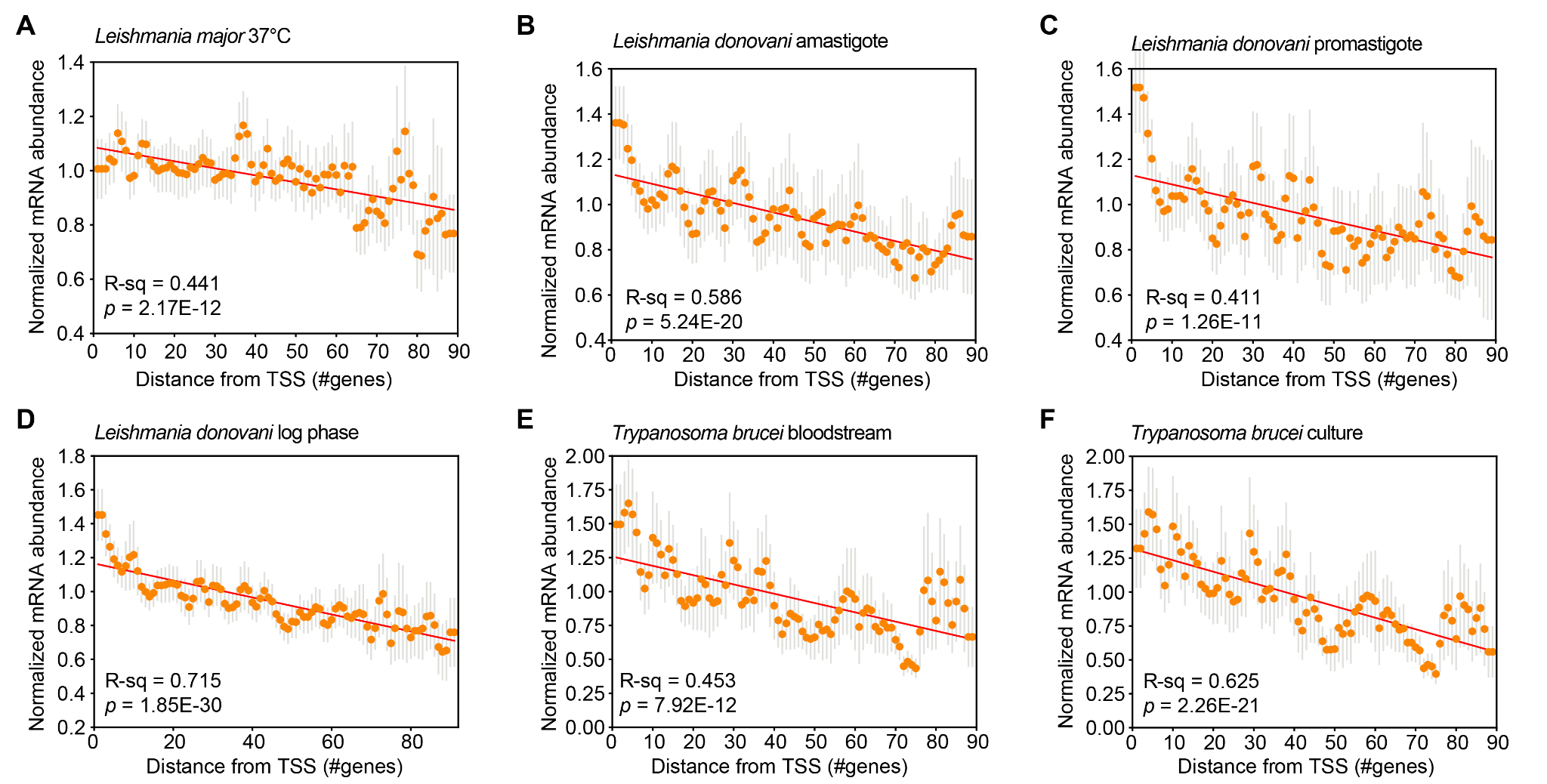

**Figure S5. Positional gradients of steady-state mRNA abundance along trypanosomatid PTUs.** (**A**) Meta-polycistron profile of steady-state mRNA abundance[[1]](https://paperpile.com/c/uXYxTn/VvMYe) for *L. major* cells grown at 37°C. Genes were aligned by their index within each PTU, relative to the first gene downstream of the transcription start site. Orange points represent the mean per-gene mRNA abundance at each gene-index position, normalized to the mean mRNA abundance of the corresponding PTU; gray vertical lines denote standard deviations. The red line represents the linear regression, with the corresponding R² and *p*-value indicated. (**B**) Same as in A, for *L. donovani* cells in the amastigote phase[[2]](https://paperpile.com/c/uXYxTn/iymuX). For this analysis, *L. major* PTU boundaries were assumed to apply to the *L. donovani* genome. (**C**) Same as in A, for *L. donovani* cells in the promastigote phase[[2]](https://paperpile.com/c/uXYxTn/iymuX). (**D**) Same as in A, for L. donovani cells in log phase[[3]](https://paperpile.com/c/uXYxTn/35Mbk). (**E**) Same as in A, for *T. brucei* bloodstream-form cells[[4]](https://paperpile.com/c/uXYxTn/tGUzb). (**F**) Same as in A, for *T. brucei* cells from adipose tissue[[5]](https://paperpile.com/c/uXYxTn/JMjwG).

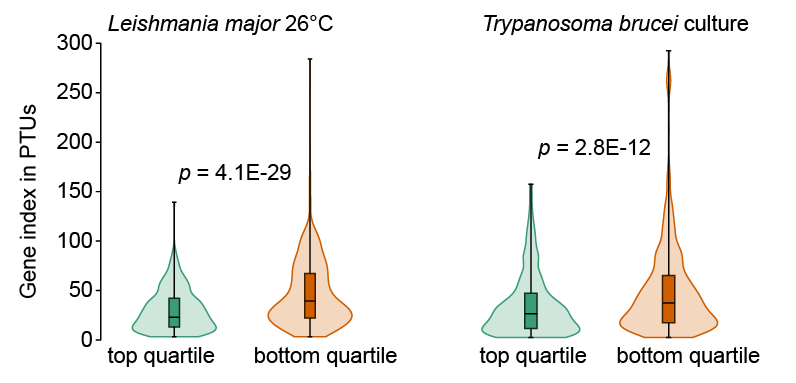

**Figure S6.** Violin plots represent gene index distributions for highly- (top quartile) and lowly- expression (bottom quartile) genes in *L. major* and *T. brucei* cells.

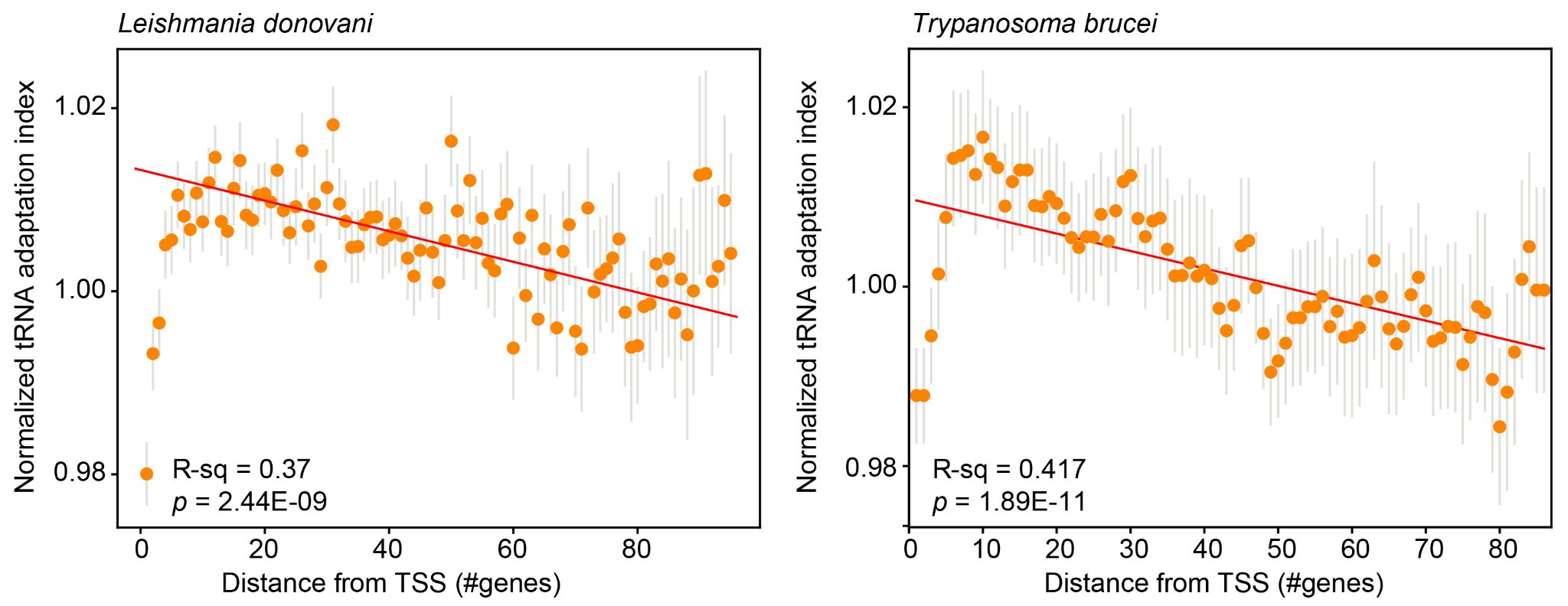

**Figure S7. Positional gradients of tRNA adaptation index along trypanosomatid PTUs.**

**Left**: Meta-polycistron profile of tRNA adaptation index (tAI) for *L. donovani* genes. Genes were aligned by their index within each PTU, relative to the first gene downstream of the transcription start site. Orange points represent the mean per-gene tAI at each gene-index position, normalized to the mean tAI of the corresponding PTU; gray vertical lines denote standard deviations. The red line represents the linear regression, with the corresponding R² and *p*-value indicated. For this analysis, *L. major* PTU boundaries were assumed to apply to the *L. donovani* genome. **Right**: Same analysis for *T. brucei* genes.

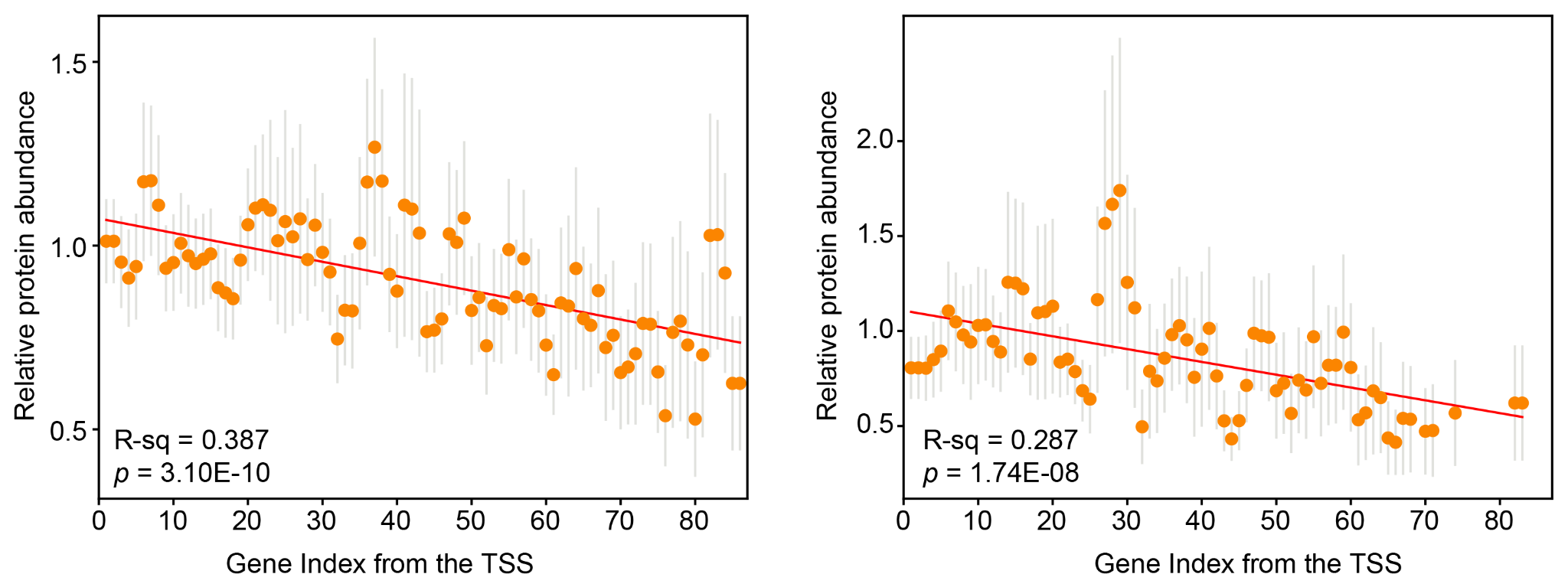

**Figure S8. Positional gradients of protein abundance along *L. major* PTUs.** Meta-polycistron profiles of protein abundance for *L. major* genes. Genes were aligned by their index within each PTU, relative to the first gene downstream of the transcription start site. Orange points represent the mean per-gene protein abundance at each gene-index position, normalized to the mean protein abundance of the corresponding PTU; gray vertical lines denote standard deviations. The red line represents the linear regression, with the corresponding R² and p-value indicated. Left and right panels correspond to proteomics datasets from Polanco et al.[[6]](https://paperpile.com/c/uXYxTn/bMtXy) and Rajan et al.[[7]](https://paperpile.com/c/uXYxTn/X5j19), respectively.

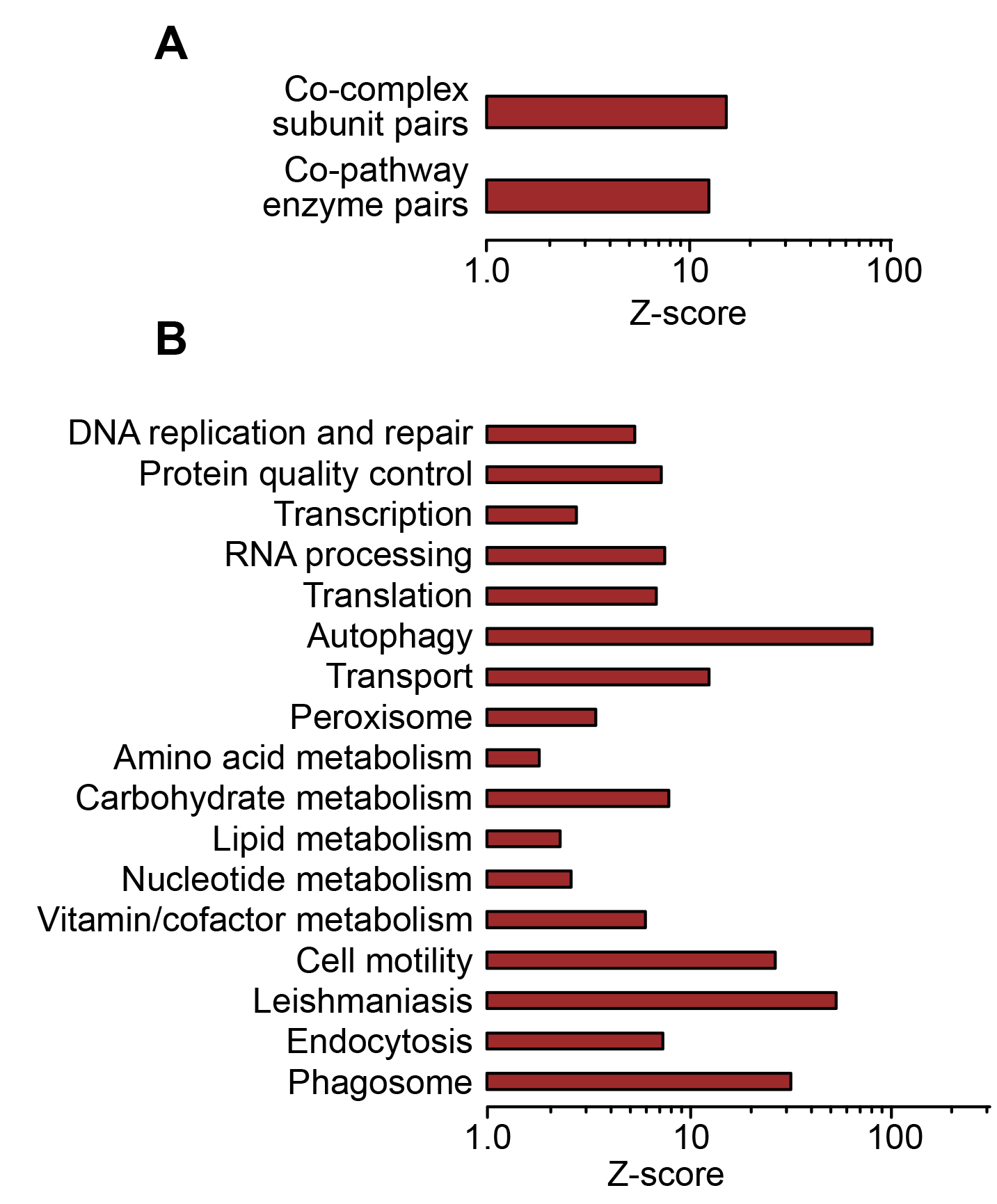

**Figure S9. Gene organization within trypanosomatid chromosomes.** (**A**) Bar plots represent enrichment of gene pairs encoding subunits of the same complex or enzymes of the same metabolic pathway within the same chromosome of *L. major*. The enrichment is quantified in terms of a Z-score against a background of 10,000 random gene pairs. (**B**) Same as **A**, for KEGG-annotated broader gene functional classes.
